# Parallel selection in domesticated Atlantic salmon from divergent founders including parallel selection on WGD-derived homeologous regions

**DOI:** 10.1101/2024.07.22.604627

**Authors:** Pauline Buso, Célian Diblasi, Domniki Manousi, Jun Soung Kwak, Arturo Vera Ponce de Leon, Kristina Stenløkk, Nicola Barson, Marie Saitou

## Abstract

Aquaculture has a considerably shorter history compared to the domestication of plants and animals. Among aquatic species, those that have undergone whole genome duplication events (WGD) seem particularly successful. This suggests that genetic redundancy from WGD is important for domestication, possibly similar to plant domestication. Atlantic salmon (*Salmo salar*), which has experienced a lineage-specific WGD, has undergone rapid domestication through intensive breeding since the 1960s.

Here, we examined the genomic responses to the domestication of Atlantic salmon, including the impacts of WGD, by comparing the whole genome sequence data of aquaculture and wild populations from two lineages: the Eastern and Western Atlantic (Western Norway and North America). Our analysis revealed shared selective sweeps on identical SNPs in major histocompatibility complex (MHC) genes across distinct aquaculture populations compared to their wild counterparts. This SNP level parallelism suggests that a combination of long-term balancing selection and recent human-induced selection has significantly shaped the evolutionary trajectory of MHC genes. In addition, we observed selective sweeps on gene pairs in the homeologous regions originating from WGD, highlighting WGD’s role in maintaining genomic variation and potentially reducing pleiotropy through sub-functionalization.

This unique type of “parallel” selection contributes to adapting to the intensive artificial conditions of aquaculture. These findings provide valuable insights into the genetic mechanisms of domestication and adaptive responses in Atlantic salmon, suggesting that the salmonid whole genome duplication has underpinned their successful rapid domestication. Our research emphasizes the importance of maintaining genetic diversity to support sustainable aquaculture practices.

## INTRODUCTION

Domestication marks one of the key shifts in human history, tracing back to the enduring bond between early hunter-gatherers and wolves over 15,000 years ago (Vigne, 2011). Domesticated plants and animals became essential to human societies for social, nutritional, symbolic, or economic purposes, reflecting the development of multicultural societies that emerged at the end of the last ice age (Larson *et al*., 2014). Using domestication and artificial selection to understand the process of evolution has a long history (e.g. Darwin, 1868). Domestication refers to a small, selected fraction from a wild population adapting to a human-controlled environment. This process can change the genetic composition and phenotypes of farmed individuals in response to both intentional and unintentional selection (Wang *et al*., 2014). Small initial founder populations result in a strong bottleneck and limited genetic diversity. Consequently, the genomic evolution of domesticated species is influenced by strong genetic drift, as well as intentional artificial selection for favorable production traits (Innan and Kim, 2004), and unintentional domestication selection resulting from adaptation to the captive environment and human contact (Mignon-Grasteau *et al*., 2005; Frantz *et al*., 2020; Huang *et al*., 2022). Thus, domesticated species carry signatures of these processes in their genomes. These genomic signatures provide valuable insights into their domestication history and make them good models for genetic studies of evolution (Wiener and Wilkinson, 2011).

Conventional investigations into the evolutionary changes after domestication relied on archaeological records (Diamond, 2002). Recent advances using ‘Omics approaches have allowed researchers to trace the geographical and temporal origins and early stages of domestication, revealing the genomic basis of favored traits and selected loci in various species (Frantz *et al*., 2020). Moreover, studies using ancient DNA have provided insights into older domestications, offering a broader understanding of the domestication process at the molecular scale (Frantz *et al*., 2020). Studies using ancient DNA have also provided insights into older domestications, offering a broader understanding of the domestication process at the molecular scale. Comparative studies between farmed strains and their wild counterparts have highlighted both the beneficial genomic regions selected for specific traits through domestication and artificial selection, as well as the unintended consequences of these processes (Glover *et al*., 2017a; Fernandez *et al*., 2021; Izawa, 2022). In addition, Genome-Wide Association Studies (GWAS) has identified the genetic bases for desirable traits in various livestock, aquacultural, and agricultural species (Han *et al*., 2018; Li *et al*., 2018; Liu *et al*., 2019; Zhang *et al*., 2019; Freebern *et al*., 2020; Horn *et al*., 2020; Sokolkova *et al*., 2020; Eltaher *et al*., 2021). This approach has facilitated artificial selection towards production goals (Hayes, Lewin and Goddard, 2013; Wang *et al*., 2020; Yáñez *et al*., 2023). Besides its practical applications in functional genomics, GWAS also offers evolutionary insights, especially for artificial selection and polygenic selection. By combining the loci identified through GWAS with evolutionary studies, researchers can narrow down which genomic region have been selected and how they relate to specific traits. This integrated approach helps to uncover the genetic mechanisms driving domestication and artificial selection.

As domestication has often occurred multiple times in the same species, it can drive parallel evolution across these independently domesticated populations under similar selection pressures. This domestication selection can provide an opportunity to study selection in animals with longer generation times that are challenging to study in the laboratory. Parallel evolution refers to the process where multiple populations develop similar phenotypic or genetic adaptations independently. The probability of parallel evolution in response to similar selection pressures is dependent on the richness of genetic variation, standing genetic variation when this occurs over short time periods (Wood, Burke and Rieseberg, 2008; Bailey *et al*., 2017). A central question in understanding parallel evolution is whether parallel selection targets the same variants, genes, or biological pathways across populations.

Evolutionary convergence, where multiple populations or species evolve similar traits independently under similar selection pressures, provides compelling insights into the repeatability of adaptation at the genetic level. For instance, high-altitude adaptation has been observed not just within independent human populations but also in domesticated animals such as dogs, cows, horses, sheep, pigs, and chickens. This high altitude adaptation often involves selection on the *epas1* gene, highlighting a shared genetic response to environmental pressures (Witt and Huerta-Sánchez, 2019). Similarly, mammals that have adapted to high-starch diets, including various domesticated animals, have an increased copy number of the amylase gene (*amy*), (Pajic *et al*., 2019). In contrast, in a comparative study between sheep and goats, researchers identified 20 selective sweeps that differentiate domestic breeds from their wild counterparts in both Capra and Ovis, involving different genes that are associated with similar phenotypic expressions (Alberto *et al*., 2018). These examples underscore the complexity of convergent evolution, where both shared and distinct genetic pathways can contribute to similar adaptive outcomes across different species.

Compared to the long history of terrestrial livestock domestication, most aquaculture species have only recently been domesticated in the early 1980s (Teletchea, 2021). This recent domestication can provide a unique opportunity to investigate the earlier stages of domestication. Unlike established terrestrial livestock species, which we can see now, aquaculture species present a varied landscape of domestication success, with some species achieving breakthroughs in captive breeding while others continue to rely heavily on wild stock imports to sustain aquaculture production. Successful domestications in aquaculture are predominantly observed in families with duplicated genomes, such as sturgeons, cyprinids (including carp), and salmonids, with the exception of cichlids, which, however, have high rates of gene duplication (Brawand *et al*., 2014). This suggests a possible parallel with plant domestication, where genomic duplication plays a crucial role (Salman-Minkov, Sabath and Mayrose, 2016; Wang *et al*., 2024). Whole genome duplication results in increased redundancy, allelic diversity, heterozygosity, and meiotic recombination, augmenting adaptive plasticity, which can enhance tolerance to new mutations (Lien *et al*., 2016; Gundappa *et al*., 2022). Whole genome duplications lead to genomic redundancy and relaxed purifying selection that can result in the accumulation of genetic diversity and increased adaptability, contributing to the species’ adaptability, including domestication success (Baduel *et al*., 2019). Duplicated genomes can also retain higher levels of standing genetic variation and increase the likelihood of parallel evolution (Ebadi *et al*., 2023). Consequently, understanding the contribution of genetic redundancy to domestication, where genome duplication results in multiple gene copies performing similar functions, is crucial to reveal the evolutionary mechanisms nurturing successful domestication in aquaculture species.

Among the aquaculture species with duplicated genomes, Atlantic salmon (*Salmo salar*), in particular, has been subject to rapid and strong artificial selection and has seen growing economic importance since the 1960s (Cross and King, 1983; Ford and Myers, 2008; Glover *et al*., 2017b). Atlantic salmon is a widely distributed anadromous species characterized by a strong tendency to reproduce in the rivers of their birth, which encourages genetic divergence among populations (Klemetsen *et al*., 2003; Lehnert *et al*., 2018). Wild populations of Atlantic salmon have been split into four main phylogeographical lineages, three in Europe (East Atlantic, Barents/White Sea, Baltic) and one in North America (West Atlantic) (Bourret *et al*., 2013; Rougemont and Bernatchez, 2018). According to the estimates by Rougemont and Bernatchez, (2018), the divergence between the continents began more than one million years ago. Thus, Atlantic salmon from each continent were strictly and largely isolated during the Quaternary Ice Age before secondary intercontinental contact were facilitated by post-glacial environmental changes conducive to long-distance gene flow (Bernatchez and Wilson, 1998; Hewitt, 2000; Lehnert *et al*., 2018; Nugent *et al*., 2024). Notably, these lineages exhibit karyotypic variation, with the North American populations displaying a polymorphism in chromosome numbers due to segregating chromosomal fusions (Brenna-Hansen *et al*., 2012). Atlantic salmon in both Europe and North America have been domesticated multiple times from different source populations. Repeated domestication and artificial selection have been directed at similar traits under human-induced environments, suggesting we may observe parallel evolution between these independently domesticated lineages. The salmonid-specific whole-genome duplication event that occurred approximately 80 million years ago (Lien *et al*., 2016) could have facilitated rapid genomic adaptations in response to domestication and artificial selection pressures.

Despite the very recent domestication, studies of Atlantic salmon have revealed rapid genetic changes in pathways for metabolism, immunity, and faster growth (Debes *et al*., 2012; Bicskei *et al*., 2014; Harvey *et al*., 2018; Jin *et al*., 2020; Bull *et al*., 2022). Moreover, signatures of domestication selection have also been observed in neurological genes (Bertolotti *et al*., 2020), and behavioral alterations, such as increased sensitivity to predation and increased tolerance to stress compared with the wild (Solberg *et al*., 2020). So far, several studies have investigated parallel selection in Atlantic salmon across different geographic lineages. López *et al*. (2019) observed parallel evolution at four genes (e.g ubiquitin-conjugating enzyme E2F putative (*ube2f*), collagen alpha-1XIII chain (*coda1*), autism susceptibility 2 protein-like (*auts2-like*), and transient receptor potential cation channel subfamily M member 3-like (*trpm3-like*) between Scottish and North American populations (i.e Canada). however, Naval-Sanchez *et al*. (2020) did not observe any overlap among selected loci between populations of European and North American origin, highlighting the need for further research to clarify the role of parallel selection in Atlantic salmon domestication.

Here, we investigated the genomic responses of independent Atlantic salmon lineages to recent domestication and artificial selection. By examining genomic divergence between wild and aquaculture populations in Western Norway and North America, we aimed to: (1) understand the evolutionary mechanisms contributing to the initial stages of domestication by identifying specific loci and biological pathways affected. (2) Evaluate the effect of salmonid-specific genome duplication (WGD) on the domestication responses of Atlantic salmon. (3) investigate parallel evolution within and between Atlantic salmon lineages from independent domestications. To achieve these objectives, we used whole genome sequencing data, along with three selection scan methods based on allele frequencies (F_ST_), site frequency spectrum (Tajima’s D), and haplotype differentiation (XP-EHH). This approach enabled us to detect potential loci under selection, including those loci shared between the two phylogeographic lineages of Atlantic salmon.

## RESULTS

### Population genetic structure

We first examined the genetic structure of the sequenced populations. We compared the genetic patterns of aquaculture samples from MOWI (Norwegian) with their wild counterparts from 17 rivers in West Norway, the region from where this breed originates, and aquaculture samples from 3 North America breeds from the Gulf of St Lawrence/Bay of Fundy/Maine region (Gaspe, St John River and Penobscot River respectively) and 8 rivers from the Gulf of St Lawrence for comparison (**Figure 1 (A) & (B)**). Principal Component Analysis (PCA) (**Figure 1 (C)**) based on 148,433 variants derived from whole genome sequencing showed four distinct clusters: (1) Norwegian aquaculture, (2) wild Norwegian, (3) North American aquaculture from the St John and Penobscot rivers and wild North American, and (4) North American aquaculture from Gaspe. The first two principal components (PCs) jointly accounted for 64.59% of the total variance, with PC1 (53.85% of variance) separating the East Atlantic (North American) and West Atlantic (West Norwegian) lineages, and PC2 (10.74%) distinguishing between wild and aquaculture populations in Norway, while we did not observe a clear separation between wild and aquaculture populations in North America.

**Figure 1.**
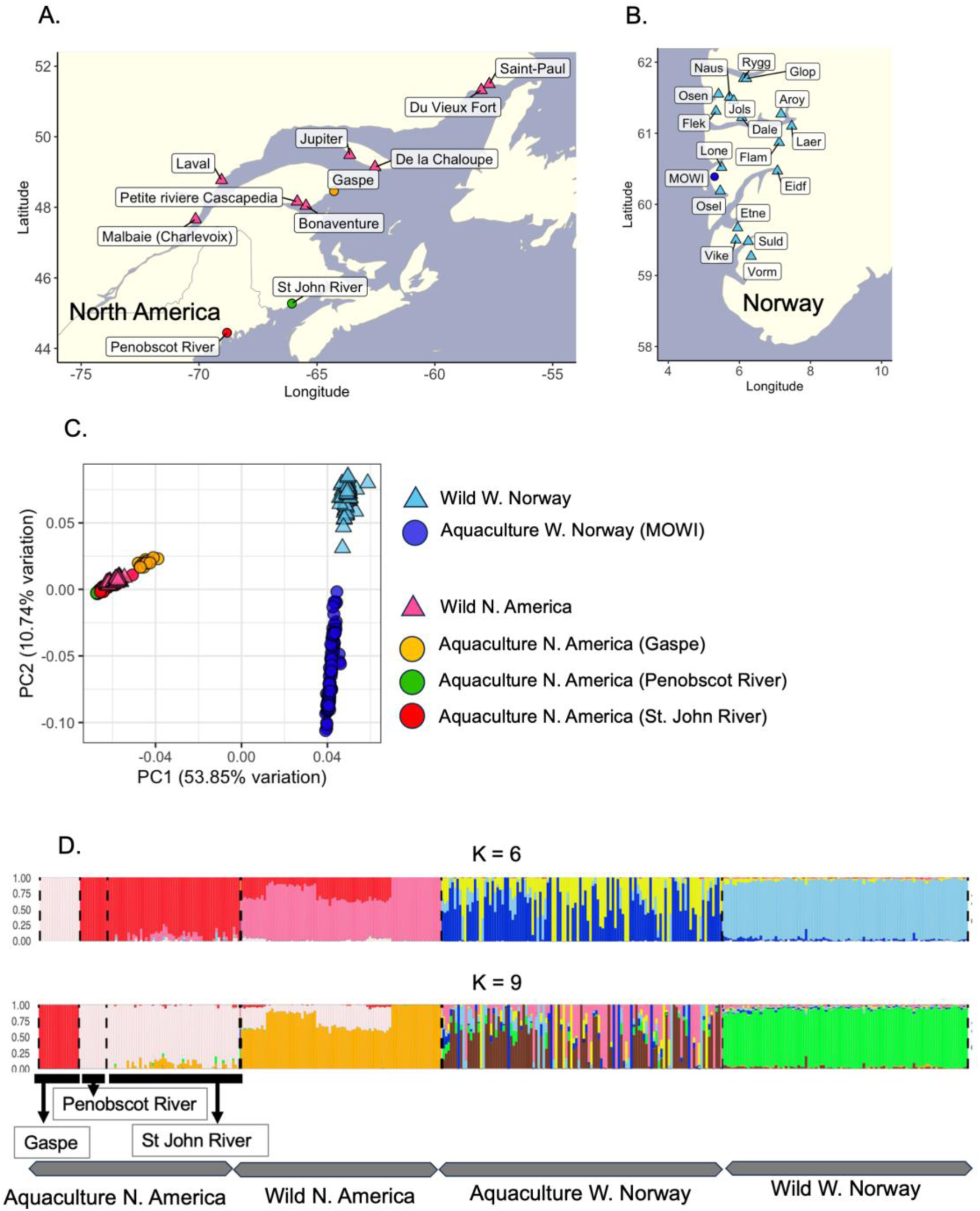
Sample distribution and genetic structure of Atlantic salmon lines from western Norway and North America. (A) Geographic distribution of Atlantic salmon population in North America. Aquaculture samples were used from Gaspe (Gaspe, n=16), St John River (n=53) and Penobscot River n=11). Wild samples came from 8 rivers: Malbaie Charlevoix (n=10), Bonaventure (n=10), Petite riviere Cascapedia (n=10), Laval (n=10), De la Chaloupe (n=10), Jupiter (n=10), Du Vieux Fort (n=10) and Saint-Paul (n=10). (B) Geographic distribution of Atlantic salmon population from Western Norway. Aquaculture samples come from MOWI Norwegian breeding line (n=112). Wild counterparts were collected among 17 ivers: Vorma (Vorm, n=2), Suldalslågen (Suld, n=10), Vikedalselva i Vindafjord (Vike, n=10), Etneelva Etne, n=2), Eidfjordvassdraget (Eidf, n=2), Oselva i Os (Osel, n=2), Loneelva i Osterøy (Lone, n=2), Flåmselva (Flam, n=10), Lærdalselva (Laer, n=2), Årøy (Aroy, n=10), Daleelva (Dale, n=2), Flekkeelva Flek, n=2), Nausta (Naus, n=10), Jølstra (Jols, n=10), Oselvvassdraget (Osen, n=10), Ryggelva (Rygg, n=10) and Gloppenelva (Glop, n=2). (C) Scatterplot of principal component analysis (PCA). Principal components 1 and 2 (PC1 and PC2) were calculated for all Atlantic salmon samples. (D) Population genetic structure analysis. All samples were clustered into 6 and 9 genetic components (K=6 and K=9) according o the lowest cross-validation error.

In agreement with the PCA, ADMIXTURE analysis (Alexander, Novembre and Lange, 2009; Alexander and Lange, 2011) also highlighted a clear separation between the two geographical regions. At K=6, where the cross-validation error plateaued (**Supplementary material, Figure S1**), each phylogeographical lineage showed distinct ancestry, as expected, with varying extents of shared ancestry among wild and aquaculture individuals from the same lineage (**Figure 1D**). Despite the recent domestication from wild populations (Gjedrem, Gjøen and Gjerde, 1991; Glover *et al*., 2009; Bradbury *et al*., 2022), the Norwegian aquaculture population (MOWI) demonstrated higher genetic heterogeneity than its wild counterpart, possibly reflecting the admixed origin from multiple wild populations. Conversely, North American aquaculture and wild populations were genetically more similar, except for the Gaspe individuals, which were largely distinct, with limited representation in wild populations. This reflects more recent co-ancestry among North American individuals resulting from the later development of aquaculture in Eastern North America. We also observed a small amount of admixture from the wild East Atlantic group in both wild and aquaculture North American populations, reflecting known past hybridization events that have occurred both through natural secondary contact and as a result of escaped aquaculture individuals with admixed European and North American ancestry (Bradbury *et al*., 2022).

Despite the distinct ancestry of the Gaspe aquaculture population, we delineated our study populations into four main groups for the following analysis: West Norwegian Aquaculture Group, Wild West Norwegian Group, North American Aquaculture Group, and Wild North American Group. This grouping was based on the tight PCA clustering of all North American populations and the smaller variance among wild Norwegian populations on PC2 than within the MOWI aquaculture strain. The groups, therefore, reflect both the continent-scale geographical origin, Western Norway versus North America, and the selective environment, those that are wild versus those that have undergone domestication.

### Identification of putatively selected regions, genes and variants

To identify potential genomic regions under artificial selection in domestic strains, we conducted a genome scan using three statistical methods: (1) population differentiation (F_ST_) (Weir and Cockerham, 1984), (2) site frequency spectrum analysis (Tajima’s D), (Tajima, 1989) and (3) cross-population extended haplotype homozygosity (XP-EHH) (Sabeti *et al*., 2002). By combining these methods, we aimed to gain sensitivity to both hard selective sweeps, where a new beneficial allele rapidly becomes dominant, reducing the local genetic diversity owing to selection for a single haplotype containing the mutation; and soft selective sweeps, where the beneficial allele arises multiple times via either recurrent mutation or exists as standing genetic variation prior to the onset of selection, and so multiple haplotypes contain the selected allele.

We calculated Tajima’s D for the two aquaculture groups separately, applying the calculation within non-overlapping 20 kbp windows. We took the top 1% Tajima’s D outliers to indicating potential directional selection. Our analysis revealed 1053 windows in the Norwegian aquaculture population and 1032 windows in the North American aquaculture populations are putatively under selection. We then calculated F_ST_ to detect genetic differentiation between the aquaculture and wild pairs in the two geographic regions. This comparison was done using non-overlapping sliding windows of 5kbp. We identified the most differentiated regions by extracting the top 1% of values as outliers. The analysis pinpointed 3,177 windows in the West Norwegian pair and 3,088 in the North American pair, respectively, as potentially under divergent selection.

Finally, we calculated XP-EHH (Cross Population Extended Haplotype Homozygosity) for each SNP, designating the wild population as the reference and the aquaculture population as the observed group in each continent. We adopted −log10(p-value) = 4 (alpha = 0.0001) as the cutoff for significant XP-EHH values (Materials and Methods). Through the XP-EHH analysis, we identified 1914 variants in Western Norway and 1982 in North America that are under putatively recent directional selection, using an FDR with alpha set at 0.0001 (**Figure 2**).

**Figure 2.**
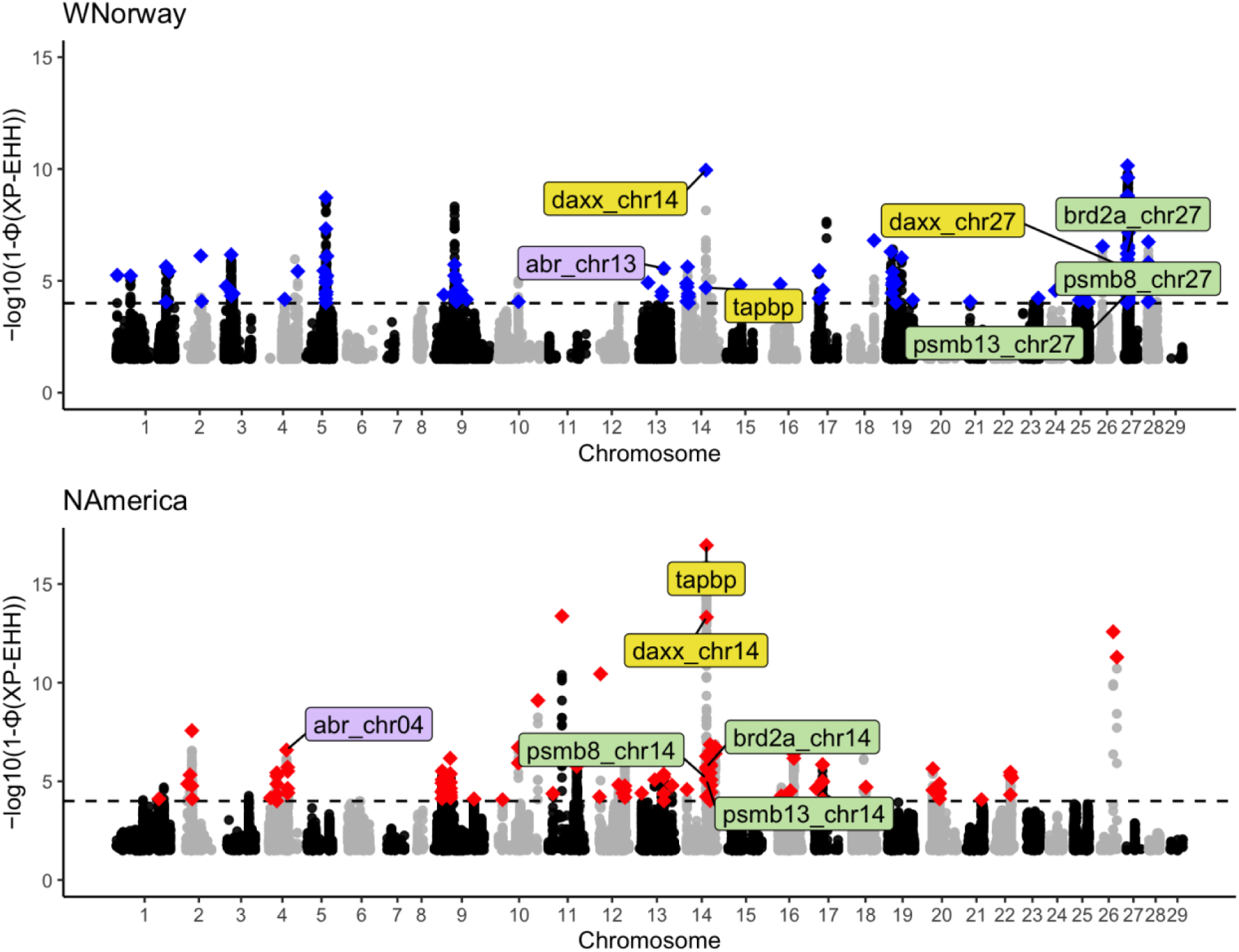
Cross-population test on Extended Haplotype Homozygosity (XP-EHH). Each dot is SNPs, while the X-axis displays the position across the genome. The Y-axis represents the −log10(p-value the XP-EHH (Cross Population Extended Haplotype Homozygosity) values. Blue Diamonds WNorway) and Red Diamonds (NAmerica) indicates SNPs with significantly high XP-EHH values that overlap genes. We adopted −log10(p-value) = 4 as the cutoff (dashed line) for significant XP-EHH values Materials and Methods). Gray and black dots are other tested SNPs. Only SNPs with a -log10(p-value) greater than 1 were plotted to ensure the computational efficiency. Labeled genes are genes with exact same SNPs under parallel selection (yellow) or selection on the homeologous regions, with matching colors n both populations showing the WGD-derived gene pairs. The suffix denotes the chromosomal location of he copy. The entire SNP list with significant XP-EHH is available in **Supplementary Table 1**.

We then asked if there were any regions or variants suggested as under selection by multiple methods, using BEDtools intersect (Quinlan, 2014). For the pairs in Western Norway and North America, we identified 54 and 9 windows, respectively, where regions were suggested to be non-neutral by both the F_ST_ and Tajima’s D metrics. Similarly, there are 144 and 363 shared SNPs suggested to be non-neutral by both F_ST_ and XP-EHH in Western Norway and North America, respectively. Meanwhile, no variants overlapped between Tajima’s D results and XP-EHH results on either continent. The limited overlap among loci detected as outliers between different methods could be because each method is sensitive to detecting selection over different timescales (Oleksyk, Smith and O’Brien, 2010). Lack of overlapped loci among methodshas been reported previously for domesticated Atlantic salmon (López *et al*., 2019).

To infer which functional groups of genes have been subjected to selection, we identified genes that overlap with windows or SNPs suggested to be non-neutral by multiple methods, using the genes from the Ensembl v111 annotation of the Atlantic salmon reference genome (version Ssal_v3.1, GCA_905237065.2). For these genes, we conducted Gene Ontology (GO) enrichment analysis using ShinyGo v0.77 (Ge, Jung and Yao, 2020). Categories were extracted where the enrichment False Discovery Rate (FDR) was < 0.05, Fold Enrichment > 2, and the Gene Ontology category contained ≥ 3 genes (**Table 1**). In the Western Norway group, genes suggested to be adaptive by both F_ST_ and Tajima’s D showed significant enrichment in the “Focal adhesion” category (FDR = 0.015). No Gene Ontology categories were detected in other groups of genes detected by multiple methods. Therefore, we expanded our investigation to include results from XPEHH alone, which can detect higher resolution and more recent directional selection. For the genes that contain SNPs identified by XPEHH, the categories “Semaphorin receptor binding (FDR = 0.01)” and “Organelle organization (FDR = 0.013)” were enriched in Western Norway, while the category “Intracellular protein-containing complex (FDR = 0.0077)” was enriched in North America. A complete list of genes overlapping with XP-EHH outliers is available in Supplementary material (**Table S1**).

**TABLE 1.**
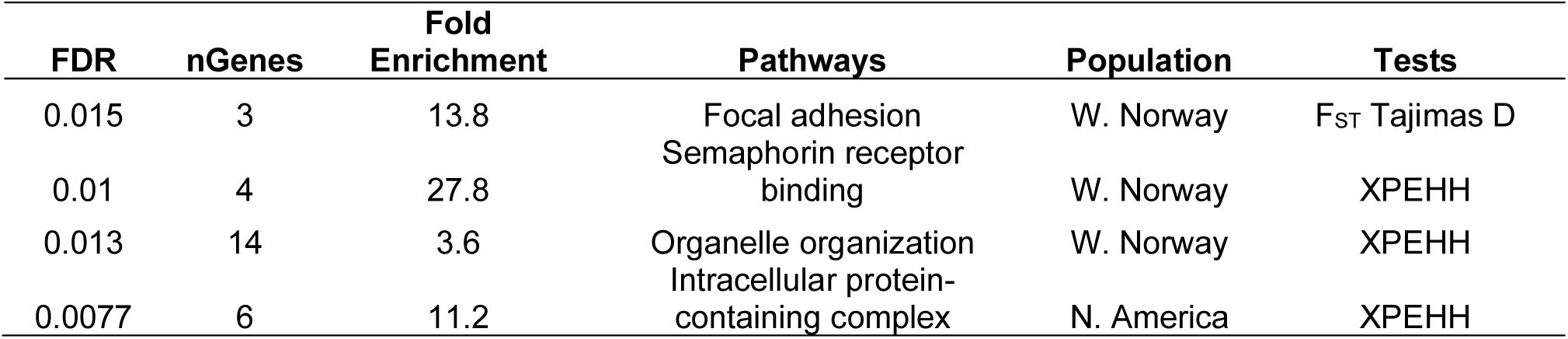
GO enrichment results on genes under putative selection. For genes that overlap with windows or SNPs suggested to be non-neutral by multiple methods, or XPEHH alone, we conducted Gene Ontology (GO) enrichment analysis using ShinyGo v0.77 (Ge, Jung and Yao, 2020). Categories are shown when the enrichment False Discovery Rate (FDR) was < 0.05, Fold Enrichment > 2, and the Gene Ontology category contained ≥ 3 our input genes (nGenes). FDR is adjusted rom the hypergeometric test. Fold Enrichment is defined as the percentage of genes in the input list belonging to a pathway, divided by the corresponding percentage in the background genes (Ge, Jung and Yao, 2020).

| FDR | nGenes | Fold<br>Enrichment | Pathways | Population | Tests |
| --- | --- | --- | --- | --- | --- |
| 0.015 | 3 | 13.8 | Focal adhesion | W. Norway | F<sub>ST</sub> Tajimas D |
| 0.01 | 4 | 27.8 | Semaphorin receptor<br>binding | W. Norway | XPEHH |
| 0.013 | 14 | 3.6 | Organelle organization | W. Norway | XPEHH |
| 0.0077 | 6 | 11.2 | Intracellular protein-containing complex | N. America | XPEHH |

### Parallel selective sweeps on the shared SNPs in the MHC locus in two Atlantic salmon lineages

We assessed parallel genomic patterns by comparing positively selected XP-EHH SNPs of aquaculture groups across continents, as this method was likely to be most specific to recent divergent selection. We found 82 SNPs with extended haplotypes in both aquaculture lineages (**Figure 3A**, **Table 1**). Of these, 58 SNPs fell within the *tapbp* gene (ENSSSAG00000040723, 14:64331687-64338464) and the rest, 24 SNPs fell within the adjacent *daxx_chr14* gene (ENSSSAG00000040701, 14:64304883-64330023) on chromosome 14. Notably, five of the shared SNPs are missense variants (**Table 2**). To check if this result reflected genotyping errors owing to the high diversity and duplicated nature of the MHC region, we manually inspected the variants using the BAM files in IGV (Robinson *et al*., 2011). We assessed that these variants are likely real (**Figure S3**) and are reliably genotyped.

**Figure 3.**
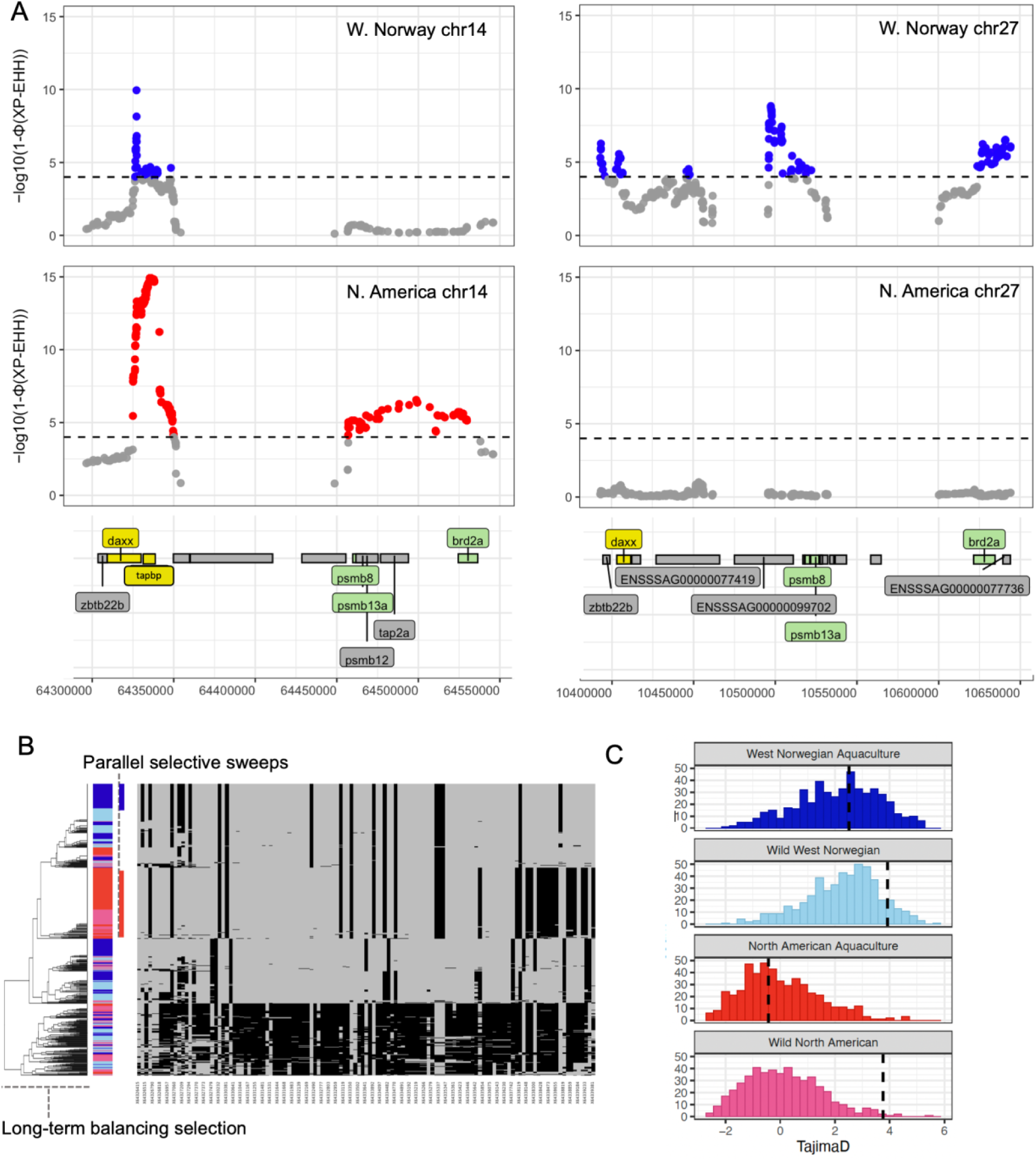
The MHC region under parallel selective sweeps in both continents A: Zoomed-in view of XP-EHH results on the MHC loci on chromosome 14 and chromosome 27 exhibiting parallel selective sweeps. X-axis displays the position across the genome. Y-axis represents the −log10(p-value the XP-EHH (Cross Population Extended Haplotype Homozygosity) values. Blue WNorway) and Red (NAmerica) indicates SNPs with significantly high XP-EHH values that overlap genes. We adopted −log10(p-value) = 4 as the cutoff (dashed line) for significant XP-EHH values (Materials and Methods). Gray and black dots are other tested SNPs. Genes with colored labels are genes under parallel selection (yellow) or parallel selection on the homeologous regions (green), with matching colors in both populations showed the WGD-derived gene pairs. Gray genes are genes with high XP-EHH values in one population. **B Haplotype pattern of the region under parallel selection.** Each column is a polymorphic site n the MHC region (126 SNPs in total, chr14: 64326415-64339381), and each row is a phased haplotype (160 North American Aquaculture (red), 224 Western Norwegian aquaculture (dark blue), 160 Wild North American pink), 196 Wild West Norwegian (light blue)). **C** Tajima’s D plots the four groups. The dashed vertical lines epresent the Tajima’s D value in the MHC locus on chromosome 14 (chr14: 64326415-64339381). The histogram represents the distribution of the Tajima’s D values from the size-matched 500 regions in each population as a neutral expectation.

**Table 2.** Outlier SNPs by the XP-EHH test, detected in both the Western Norwegian aquaculture group and North American aquaculture group. Only SNPs overlapped with genic regions in the Ensembl Varian Effect Predictor (McLaren *et al*., 2016) are shown.

| Position<br>(chr14) | Ref<br>allele | Alt<br>allele | Frequency |  |  |  | gene | variant_type |
| --- | --- | --- | --- | --- | --- | --- | --- | --- |
|  |  |  | Aqua_<br>WNorway | Aqua_<br>NAmerica | Wild_<br>WNorway | Wild_<br>NAmerica |  |  |
| 64326415 | C | T | 0.02 | 0.03 | 0.15 | 0.09 | daxx | up/downstream |
| 64326484 | G | A | 0.42 | 0.02 | 0.35 | 0.04 | daxx | up/downstream |
| 64326515 | A | C | 0.18 | 0.10 | 0.38 | 0.40 | daxx | up/downstream |
| 64326609 | T | G | 0.43 | 0.78 | 0.34 | 0.49 | daxx | up/downstream |
| 64326790 | A | T | 0.02 | 0.05 | 0.16 | 0.11 | daxx | up/downstream |
| 64326801 | C | T | 0.02 | 0.05 | 0.15 | 0.11 | daxx | up/downstream |
| 64326818 | C | G | 0.17 | 0.09 | 0.39 | 0.42 | daxx | up/downstream |
| 64326952 | T | G | 0.19 | 0.06 | 0.38 | 0.38 | daxx | up/downstream |
| 64326957 | C | G | 0.18 | 0.07 | 0.37 | 0.36 | daxx | up/downstream |
| 64327057 | A | C | 0.71 | 0.84 | 0.88 | 0.77 | daxx | up/downstream |
| 64327060 | T | A | 0.15 | 0.05 | 0.14 | 0.35 | daxx | up/downstream |
| 64327182 | T | G | 0.53 | 0.18 | 0.60 | 0.45 | daxx | up/downstream |
| 64327209 | T | C | 0.58 | 0.10 | 0.67 | 0.43 | daxx | up/downstream |
| 64327290 | A | C | 0.27 | 0.05 | 0.28 | 0.30 | daxx | up/downstream |
| 64327294 | C | A | 0.45 | 0.14 | 0.34 | 0.17 | daxx | up/downstream |
| 64327355 | C | G | 0.26 | 0.10 | 0.42 | 0.40 | daxx | up/downstream |
| 64327370 | C | T | 0.14 | 0.05 | 0.21 | 0.34 | daxx | up/downstream |
| 64327372 | T | C | 0.14 | 0.05 | 0.21 | 0.33 | daxx | up/downstream |
| 64327373 | G | A | 0.14 | 0.05 | 0.21 | 0.32 | daxx | up/downstream |
| 64327387 | C | T | 0.13 | 0.04 | 0.22 | 0.33 | daxx | up/downstream |
| 64327479 | G | A | 0.45 | 0.04 | 0.37 | 0.16 | daxx | up/downstream |
| 64330356 | A | G | 0.15 | 0.10 | 0.22 | 0.32 | daxx | up/downstream |
| 64330381 | G | C | 0.55 | 0.97 | 0.55 | 0.84 | daxx | up/downstream |
| 64330609 | G | A | 0.45 | 0.03 | 0.42 | 0.16 | daxx | up/downstream |
| 64332437 | T | A | 0.46 | 0.13 | 0.40 | 0.11 | tapasin | missense_variant |
| 64332490 | A | G | 0.16 | 0.07 | 0.30 | 0.41 | tapasin | up/downstream |
| 64332602 | C | G | 0.09 | 0.01 | 0.14 | 0.23 | tapasin | missense_variant |
| 64332777 | C | G | 0.18 | 0.08 | 0.31 | 0.43 | tapasin | intron_variant |
| 64332793 | G | T | 0.17 | 0.09 | 0.30 | 0.42 | tapasin | intron_variant |
| 64332803 | C | T | 0.18 | 0.08 | 0.31 | 0.42 | tapasin | intron_variant |
| 64332931 | T | G | 0.18 | 0.09 | 0.31 | 0.39 | tapasin | intron_variant |
| 64332959 | A | G | 0.18 | 0.08 | 0.31 | 0.40 | tapasin | intron_variant |
| 64333061 | C | G | 0.55 | 0.97 | 0.58 | 0.84 | tapasin | missense_variant |
| 64333119 | C | T | 0.17 | 0.09 | 0.30 | 0.42 | tapasin | up/downstream |
| 64333218 | A | G | 0.17 | 0.11 | 0.31 | 0.42 | tapasin | up/downstream |
| 64333350 | G | C | 0.46 | 0.91 | 0.39 | 0.56 | tapasin | splice_variant |
| 64333469 | T | C | 0.02 | 0.03 | 0.12 | 0.03 | tapasin | intron_variant |
| 64333502 | G | C | 0.15 | 0.04 | 0.24 | 0.37 | tapasin | intron_variant |
| 64333588 | C | T | 0.17 | 0.07 | 0.33 | 0.43 | tapasin | intron_variant |
| 64333641 | G | A | 0.28 | 0.03 | 0.19 | 0.16 | tapasin | missense_variant |
| 64333649 | C | A | 0.18 | 0.10 | 0.31 | 0.42 | tapasin | up/downstream |
| 64333892 | G | C | 0.46 | 0.92 | 0.34 | 0.54 | tapasin | intron_variant |
| 64334015 | T | C | 0.18 | 0.09 | 0.30 | 0.40 | tapasin | intron_variant |
| 64334097 | A | T | 0.18 | 0.10 | 0.31 | 0.40 | tapasin | intron_variant |
| 64334205 | T | C | 0.76 | 0.97 | 0.92 | 0.83 | tapasin | intron_variant |
| 64334482 | A | C | 0.55 | 0.10 | 0.62 | 0.38 | tapasin | intron_variant |
| 64334727 | C | A | 0.12 | 0.08 | 0.24 | 0.41 | tapasin | intron_variant |
| 64334770 | C | G | 0.76 | 0.97 | 0.93 | 0.85 | tapasin | intron_variant |
| 64334795 | T | C | 0.17 | 0.09 | 0.24 | 0.39 | tapasin | intron_variant |
| 64334891 | C | T | 0.14 | 0.09 | 0.28 | 0.38 | tapasin | intron_variant |
| 64335174 | T | G | 0.17 | 0.07 | 0.31 | 0.38 | tapasin | intron_variant |
| 64335192 | C | T | 0.17 | 0.08 | 0.30 | 0.37 | tapasin | intron_variant |
| 64335203 | T | C | 0.16 | 0.07 | 0.30 | 0.37 | tapasin | intron_variant |
| 64335219 | A | G | 0.16 | 0.08 | 0.30 | 0.38 | tapasin | intron_variant |
| 64335265 | G | A | 0.17 | 0.09 | 0.28 | 0.39 | tapasin | intron_variant |
| 64335266 | A | C | 0.17 | 0.09 | 0.28 | 0.39 | tapasin | intron_variant |
| 64335275 | G | C | 0.07 | 0.05 | 0.08 | 0.16 | tapasin | intron_variant |
| 64335279 | T | C | 0.16 | 0.10 | 0.27 | 0.37 | tapasin | intron_variant |
| 64335336 | T | C | 0.46 | 0.91 | 0.37 | 0.57 | tapasin | intron_variant |
| 64335337 | G | A | 0.46 | 0.92 | 0.37 | 0.57 | tapasin | intron_variant |
| 64335342 | G | C | 0.46 | 0.91 | 0.37 | 0.58 | tapasin | intron_variant |
| 64335347 | T | C | 0.17 | 0.09 | 0.25 | 0.40 | tapasin | intron_variant |
| 64335348 | G | T | 0.17 | 0.09 | 0.25 | 0.40 | tapasin | intron_variant |
| 64335361 | G | A | 0.17 | 0.05 | 0.25 | 0.25 | tapasin | intron_variant |
| 64335405 | T | G | 0.17 | 0.08 | 0.26 | 0.39 | tapasin | intron_variant |
| 64335423 | A | T | 0.17 | 0.09 | 0.26 | 0.39 | tapasin | intron_variant |
| 64335441 | G | T | 0.17 | 0.08 | 0.25 | 0.40 | tapasin | intron_variant |
| 64335446 | C | A | 0.17 | 0.09 | 0.25 | 0.40 | tapasin | intron_variant |
| 64335592 | T | C | 0.17 | 0.08 | 0.28 | 0.40 | tapasin | intron_variant |
| 64335642 | A | C | 0.29 | 0.07 | 0.42 | 0.40 | tapasin | intron_variant |
| 64335771 | A | G | 0.23 | 0.80 | 0.36 | 0.49 | tapasin | missense_variant |
| 64338300 | A | C | 0.25 | 0.03 | 0.09 | 0.16 | tapasin | up/downstream |
| 64338759 | T | A | 0.33 | 0.02 | 0.33 | 0.03 | tapasin | up/downstream |
| 64338819 | A | T | 0.42 | 0.03 | 0.35 | 0.16 | tapasin | up/downstream |
| 64338852 | C | A | 0.17 | 0.81 | 0.32 | 0.73 | tapasin | up/downstream |
| 64338859 | G | C | 0.17 | 0.81 | 0.30 | 0.73 | tapasin | up/downstream |
| 64339170 | C | T | 0.15 | 0.81 | 0.24 | 0.74 | tapasin | up/downstream |
| 64339184 | G | A | 0.15 | 0.04 | 0.24 | 0.37 | tapasin | up/downstream |
| 64339220 | A | T | 0.15 | 0.80 | 0.24 | 0.75 | tapasin | up/downstream |
| 64339233 | C | T | 0.49 | 0.82 | 0.49 | 0.90 | tapasin | up/downstream |
| 64339330 | C | A | 0.43 | 0.03 | 0.33 | 0.15 | tapasin | up/downstream |
| 64339381 | T | C | 0.11 | 0.78 | 0.20 | 0.59 | tapasin | up/downstream |

Variation in the MHC region is known to be maintained by long-term parallel selection as evidenced by trans-species polymorphisms (Tsukamoto *et al*., 2012; Teixeira *et al*., 2015). To investigate its evolutionary history, we conducted haplotype analysis and calculated Tajima’s D on this MHC region on chromosome 14 and random control regions in the four groups. Haplotype clustering using the 125 MHC SNPs on chromosome 14 (chr14:64326415-64339381) suggested that there are two independent selective sweeps in the West Norwegian aquaculture group and North American Aquaculture group and that high diversity in the haplotypes has been maintained in Atlantic salmon since before the lineage divergence between the lineages of North America and Europe (**Figure 3B**). We observed high Tajima’s D values in this region, size-matching 500 control regions, suggesting that a high level of variation has been preserved in Atlantic salmon since before the lineage divergence between North America and Western Norway. In the North American aquaculture group the Tajima’s D value for this region was in the top 56% when compared to the control regions. However, in the wild populations, the Tajima’s D value for this region was situated at the 1st percentile in comparison to the control regions. In contrast, in Western Norway, the Tajima’s D value for this region was at the 43rd percentile when compared to the control regions. In the aquaculture group. In the wild populations, the Tajima’s D value for this region was situated at the 10 percentile in comparison to the control regions (**Figure 3 C**). These results suggest that the wild group in North America shows signs of long-term balancing selection in the MHC region on chr14, while the aquaculture populations appear to have undergone selective sweeps since domestication began. Similar trends were observed in the Western Norwegian groups; however, likely due to demographic influences, the distribution of Tajima’s D values in the control regions is generally higher in Western Norway compared to those in North America (**Figure 3 C**).

### Parallel signatures of selection in alternate ohnologue copies across lineages

To further investigate how the salmonid specific WGD contributes to domestication of Atlantic salmon genetically, we examined whether one of the gene pairs from salmonid-specific WGD has adaptive signatures in the Western Norway aquaculture group while the other gene pair has adaptive signatures in the North American aquaculture group. By investigating XP-EHH outlier SNPs in genic regions, we detected three loci with parallel ohnologue aquaculture-specific selective sweeps, one containing potentially multiple sweep signals. Remarkably, in addition to the *daxx* gene on chromosome 14, which was identified as a gene under selection in both lineages, a WGD-originated duplicate of the *daxx* gene on chromosome 27 (ENSSSAG00000077402) was identified as under selection in the Western Norwegian Aquaculture Group. In addition, we identified three ohnologue gene pairs under parallel domestication-related sweeps across the two distinct geographical regions: *brd2 and psmb13* within 100kb of *daxx* on chromosomes 14 and 27, and *abr* (chromosomes 4 and 13. For these genes, we found selection signatures on one copy in the Western Norwegian group and selection signatures on the other copy of the same genes in the North American group (**Table 2)**. Such selection signatures on WGD-derived gene duplicates imply that the salmonid-specific WGD event has contributed to successful salmon domestication, highlighting the evolutionary role of elevated genetic redundancy following whole genome duplication (Gundappa *et al*., 2022).

In the Western Norwegian group, selection signatures were detected on the bromodomain-containing 2a (brd2a) gene, which is located in the MHC region on chromosome 27 (*brd_chr27*), while in the North American group, the other copy, *brd*_chr14, was also found to be under selection. Notably, the *brd2* gene has previously been reported as under positive selection in Atlantic salmon in aquaculture in Canada compared to their wild counterparts (López *et al*., 2019). Similarly, another MHC gene, proteasome 20S subunit beta 13a (*psmb13a) gene*, showed parallel selection signatures. In our analysis, selection signatures were detected on the *psmb13a_chr27* in the Western Norwegian group, and *psmb13_chr14* in the North American group.

In addition to the shared genes above, the MHC class I region also contains salmonid-lineage-specific gene copies with overlapping XP-EHH outliers, *uba, vhsv, col11A2, tcf19, zbtb22b, psmb8a* and vsp52 on chromosome 27 in Western Norwegian aquaculture. In contrast, *psmb8b* and tpa2 genes on the homeolog chromosome 14 in North American aquaculture showed the selection signatures, as reported in previous studies (Lukacs *et al*., 2007, 2010) (**Table 3**). These lineage-specific selection responses on homeolog regions indicate parallel selection targeting the MHC class 1 region.

**Table 3.** Genes under parallel selection based on the XP-EHH test. We described the gene name in this table if the gene or duplicated gene pair is overlapped with one or more outlier SNP(s) in XP-EHH test in the Western Norweigian aquaculture group and North American aquaculture group. Start and End refer to the starting and ending locations of genes on the chromosome, respectively, based on the Ssa v.3.1 annotation on Ensembl. Duplicated genes, gray shade, (**See also Figure 2 and Figure 3**) are defined according to Gillard *et al*., 2021.

| Gene Ensembl ID | Gene name | XP-EHH signatures | Region | Chr | Start | End |
| --- | --- | --- | --- | --- | --- | --- |
| ENSSSAG00000040701 | daxx_chr14 | Both_Populations | MHC | 14 | 64,304,883 | 64,330,023 |
| ENSSSAG00000040723 | tapbp | Both_Populations | MHC | 14 | 64,331,687 | 64,338,464 |
| ENSSSAG00000093386 | tap2a | NAmerica | MHC | 14 | 64,476,753 | 64,493,938 |
| ENSSSAG00000085704 | psmb13a_chr14 | Parallel_NAmerica | MHC | 14 | 64,463,937 | 64,467,854 |
| ENSSSAG00000041842 | psmb8b_chr14 | Parallel_NAmerica | MHC | 14 | 64,459,783 | 64,462,906 |
| ENSSSAG00000042614 | brd2a_chr14 | Parallel_WNorway | MHC | 14 | 64,524,519 | 64,536,467 |
| ENSSSAG00000077402 | daxx_chr27 | Parallel_WNorway | MHC | 27 | 10,402,918 | 10,411,454 |
| ENSSSAG00000077187 | tcf19 | WNorway | MHC | 27 | 10,323,665 | 10,328,897 |
| ENSSSAG00000077419 | uba | WNorway | MHC | 27 | 10,426,981 | 10,465,906 |
| ENSSSAG00000078726 | vhsv | WNorway | MHC | 27 | 10,801,247 | 10,804,941 |
| ENSSSAG00000078638 | vps52 | WNorway | MHC | 27 | 10,789,984 | 10,840,875 |
| ENSSSAG00000077397 | zbtb22b | WNorway | MHC | 27 | 10,394,411 | 10,398,690 |
| ENSSSAG00000077772 | col11a2 | WNorway | MHC | 27 | 10,645,732 | 10,694,689 |
| ENSSSAG00000077561 | psmb13a_chr27 | Parallel_WNorway | MHC | 27 | 10,521,614 | 10,527,869 |
| ENSSSAG00000085574 | psmb8a_chr27 | Parallel_WNorway | MHC | 27 | 10,518,241 | 10,521,084 |
| ENSSSAG00000077713 | brd2a_chr27 | Parallel_NAmerica | MHC | 27 | 10,621,388 | 10,634,539 |
| ENSSSAG00000048882 | abr_chr13 | Parallel_WNorway |  | 13 | 77,743,389 | 77,966,280 |
| ENSSSAG00000001855 | abr_chr4 | Parallel_NAmerica |  | 4 | 56,687,253 | 56,919,875 |

The other region with parallel selection signature is on a different homeologue region of chromosome 4, which is a duplicate of part of chromosome 13. We observed that *abr* (ABR activator of RhoGEF and GTPase) gene copy on chromosome 13 in the Western Norwegian group and the duplicate *abr_chr4* in the North American group are under selection (**Figure2, Table3**).

We observed two regions with parallel selection on salmon-specific WGD-derived ohnologues, and one major region is the MHC region with high Tajima’s D value in the aquaculture populations. This suggests balancing selection has been more important in maintaining adaptive immune variation across duplicates than prolonged tetraploid recombination, as experienced in the late rediploidization LORe regions (Robertson *et al*., 2017). Despite the occurrence of parallel ohnologue selection, the majority of selection signatures are asymmetric on gene duplicates, including in LORe regions where divergence and, therefore, the potential for functional divergence is limited, similar to the observation in the common carp genome (Wang *et al*., 2024). This suggests the major advantage of polyploidy in aquaculture is derived from reduced pleiotropy after the WGD (Marchant *et al*., 2019).

### Expression pattern of the putatively selected genes

We leveraged published RNA-seq datasets spanning various developmental stages and tissues. This allowed us to infer the functions of these putatively adaptive genes and identify potential functional divergence of homeolog gene copies. We used RNA sequencing data from early development stages (AQUA-FAANG project No. PRJEB51855) and from various tissues of healthy fish (NCBI Project No. SRP011583) of European Atlantic salmon (**Materials and Methods**).

Both daxx homeologue gene copies, located on chromosomes 14 and 27, were under selection. The daxx_chr14 shows signatures of selective sweep in both the North American and West Norwegian aquaculture lineages, whereas the daxx_chr27 exhibits signs of selective sweep only in the West Norwegian aquaculture lineage. Both copies showed high expression in the ovary, testis, and during early blastulation stages (**Figure 4**), suggesting a role in both reproduction and early development. Additionally, daxx_chr27 is also highly expressed in the gill, spleen, and head kidney, indicating its role in immunity. This pattern suggests some retained shared function and possible functional divergence of the duplicated gene copies. Increased *daxx* gene expression has been related to high stocking density in rainbow trout, potentially related to the anti-infection mechanism (Détrée and Gonçalves, 2019). In addition to the immunological role, *daxx* is also known to play a key role in apoptosis in various species (Khelifi, D’Alcontres and Salomoni, 2005; Yao *et al*., 2014).

**Figure 4.**
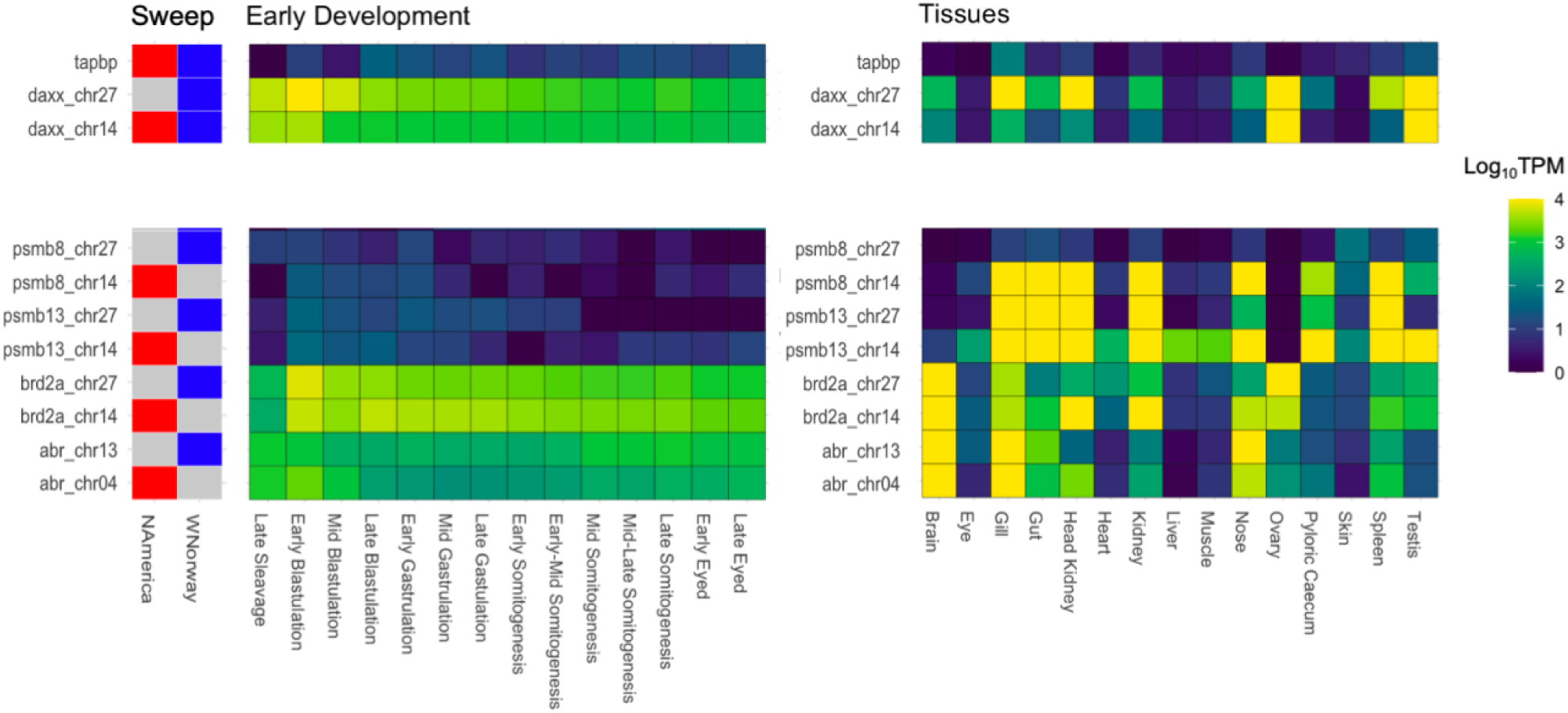
Expression of genes under parallel selection or parallel selection on the homeologous regions in early development stages and mature tissues. Gene expression is represented as log-transformed Transcripts Per Million (TPM). The sweep panel indicates the gene that have undergone selective sweeps in North America (red) or Western Norway (blue). The left heatmap shows the gene expression levels during early development, from late gastrulation to the eyed egg stage. The right heatmap shows the gene expression levels across different tissues in the mature Atlantic salmon. We used publicly available RNA sequencing data from early development stages (AQUA-FAANG project No. PRJEB51855) and from various tissues of healthy fish (NCBI Project No. SRP011583) including brain, eye, gill, gut, head kidney, heart, kidney, liver, muscle, nose, ovary, caecum, skin, spleen and testis of Atlantic salmon in Europe.

The *tapbt* gene, located adjacent to the *daxx* gene in the MHC region on chromosome 14 and showing selection signatures in both populations, showed low expression in most tissues in the healthy fish. This gene was, though, moderately expressed in the gill (**Figure 4**), which acts as a gateway for potential pathogens, highlighting its potential role in the immune response (Amill *et al*., 2024). It is reported that *tapbt* expression is upregulated in a rainbow trout monocyte/macrophage cell line in response to viral infection (Sever *et al*., 2014).

Both copies of *brd2* show high expression in the brain and gill, while *brd_chr27* is also expressed in ovary and *brd_chr14* is expressed in head kidney, kidney and nose (**Figure 4**). As with daxx, this pattern indicates that each copy may have unique as well as overlapping roles in different tissues. Both copies also display high expression levels during the early stages of development, particularly in early blastulation (**Figure 4**). This is consistent with the established roles of *brd2* in reproductive and developmental processes, including spermatogenesis and folliculogenesis (Rhee *et al*., 1998), and egg polarity in zebrafish (*Danio rerio*) (DiBenedetto *et al*. 2008). *Brd2* is also responsible for nervous system development and morphogenesis and differentiation of the digestive tract in zebrafish (Murphy *et al*. 2017). In the Gene Ontology analysis, both *daxx* and *brd2* appear in the organelle organization category that is enriched in genes under selection in Norwegian aquaculture (**Table 1**). Another duplicate pair under parallel selection *Psmb8,* under aquaculture selection on chromosomes 27 and 14 in Norwegian and North American, respectively, codes for a beta subunit of the type 8 proteasome, which facilitates endopeptidase activity and proteasomal protein catabolism (Arima *et al*., 2011). *Psmb8a_chr14* is highly expressed in many tissues, including immune-related tissues such as the gill, head kidney, and spleen, while *psmb8a_chr27* did not show such high expression, suggesting possible functional divergence (**Figure 4**).

*Abr*, on chromosomes 13 and 4 and under homeologous parallel selection in Norwegian and North American aquaculture, respectively (**Table 3**), is an activator of RhoGEF and GTPase, which activate cerebellar development in multiple vertebrates (Chuang *et al*., 1995; Mulherkar *et al*., 2014; Guo, Yang and Shi, 2020). Both copies of *abr* were highly expressed in the brain (**Figure 4**), suggesting a neurological function in Atlantic salmon as well.

## DISCUSSION

The origins of animal and plan domestication trace back around 12,000 years ago, evolving through complex interactions among humans, animals and plants shaped by ecological, biological, and cultural forces (Larson *et al*., 2014). During this process, humans managed the reproduction of livestock and crops, selectively breeding for traits that enhanced survival and productivity. In contrast, aquaculture generally has a much shorter history, though the domestication of common carp in China dates back 8,000 years to (Nakajima *et al*., 2019). The sector only boomed in the early 1980s, becoming the fastest-growing area of food production despite numerous challenges from biological and technical gaps (Teletchea, 2021). Notably, the families Cyprinidae, Salmonidae, and Acipenseridae have seen significant success (Teletchea, 2021). Atlantic salmon, in particular, has emerged as a prime example of rapid domestication due to systematic breeding efforts in recent decades (Houston and Macqueen, 2019).

### Parallel selective sweeps in MHC genes in aquaculture across continents: Likely retention of variation under long-term balancing selection

In our study, we investigated the genomic responses of Atlantic salmon populations to recent domestication by conducting selection scans between wild and aquaculture population pairs in Western Norway and North America. Our comparative study identified 82 SNPs within the *daxx* and adjacent *tapbt* genes, including five nonsynonymous SNPs, shared selective sweeps in aquaculture population pairs in Western Norway and North America. These genes play crucial roles in the immune system: *daxx* and *tapbt* are both members of the major histocompatibility complex (MHC) class IA region in Atlantic salmon (Grimholt, 2018). *Tapbt* encodes a transmembrane glycoprotein, which stabilizes the peptide loading complex of MHC class I molecules, aiding in T cell immune recognition (Ortmann *et al*., 1997; Lehner, Surman and Cresswell, 1998; Garbi *et al*., 2003). In addition to the immunological function, *daxx* also plays a key role in apoptosis (Khelifi, D’Alcontres and Salomoni, 2005; Yao *et al*., 2014; Détrée and Gonçalves, 2019).

Our findings suggest parallel selective sweeps predominantly at the MHC regions in distinct aquaculture groups, following long-term balancing selection and continental population divergence (around 1M years) (Rougemont and Bernatchez, 2018). This result implies that non-MHC genomic regions hold too few variants shared across continents for parallel responses at the SNP level, likely due to fixation by genetic drift. In contrast, variants in the MHC regions have been preserved due to long-term balancing selection. This preservation allows room for parallel selection on the shared variants between continents.

In the MHC region, which is thought to be under negative frequency-dependent selection, extreme long-term balancing selection is observed in vertebrates (McConnell *et al*., 2016; Veríssimo *et al*., 2023). Likewise, in natural environments, we can assume that the allelic diversity at the MHC locus of Atlantic salmon has been maintained through balancing selection. However, in aquaculture settings in both Western Norway and North America, specific alleles have been selected due to common selective pressures for increased immune response to common problem diseases in high-density net pen populations and associated elevated disease prevalence (Krkosek, 2010). Alternatively, this selection may target phenotypic traits linked to other functional effects of these genes, exemplified by the fact that *daxx* also plays a key role in apoptosis (Michaelson *et al*., 1999; Khelifi, D’Alcontres and Salomoni, 2005). While nonsynonymous SNPs have been identified, pinpointing the causal variants needs more targeted approaches, such as MHC-specific sequencing, to fully resolve this rapidly evolving region (Sundaram *et al*., 2020) and functional analysis, such as gene editing. In the long term, co-evolution between hosts and pathogens could lead to currently advantageous alleles becoming disadvantageous (Spurgin and Richardson, 2010). Maintaining genetic diversity will likely contribute to sustainable aquaculture practices.

### Parallel selection on homeologous regions: genome duplication as a possible driver of successful Salmonid aquaculture

Many aquaculture trials for new species fail within a few years, due to the challenges including: (1) developing suitable feeds for the initial feeding of fish larvae, (2) poor gonadal development, and (3) lack of captive spawning (Teletchea, 2021). Despite these challenges, the families Cyprinidae, Salmonidae, and Acipenseridae have successfully overcome these bottlenecks. A common trait among these families is their lineage-specific whole genome duplication (WGD), which may provide a genomic potential and resources enough in the populations to thrive in the novel, artificial environment and intensive selection of aquaculture (Lien *et al*., 2016; Chen *et al*., 2019; Xu *et al*., 2019; Du *et al*., 2020; Redmond *et al*., 2023; Rondeau *et al*., 2023). Many plant species with duplicated genomes have also been successfully cultivated, such as wheat, maize, potato, and brassicas (The Brassica rapa Genome Sequencing Project Consortium *et al*., 2011; Salman-Minkov, Sabath and Mayrose, 2016; Zhang, Wang and Cheng, 2019). This suggests that WGD can increase the tolerance for various environmental conditions, including domestic environments, by offering sufficient genetic variation plasticity.

Supporting the concept of WGD as a potential facilitator of rapid adaptation and therefore enhanced resilience in aquaculture, we identified four WGD-originated gene pairs under parallel domestication-related selection across two distinct geographical lineages: psmb8a, psmb13a, brd2a on chr14 and chr27 (genes on MHC regions), and abr on chr4 and chr13 (genes involved in neurological functions). This occurrence is notably rare, likely due to the ancient nature of the WGD event, resulting in the diversification or pseudogenization. We specifically highlight the MHC gene pairs (chr14 and chr27) as they exhibit parallel selection, where variation may have been maintained over long periods by frequency-dependent selection. These genes are involved in immune, neurological, and early developmental processes with some pleiotropic effects, each playing a crucial role in the adaptation of Atlantic salmon to aquaculture environments. For these genes, we found signatures of recent selective sweeps on one copy in the Western Norwegian group and selection signatures on a duplicate copy of the same genes on another chromosome in the North American group. This phenomenon can be characterized as a unique type of “parallel” selection specific to species that have undergone genome duplication and domestication. In this context, the contribution of WGD seems most likely to be: (1) allowing for increased immune variation maintained among duplicated immune gene complexes and (2) reducing pleiotropy through sub-functionalization (Marchant *et al*., 2019). Similar patterns have been reported in traits such as vibrant skin color in common carp and leaf-heading traits in Brassica species (Cheng *et al*., 2016; Wang *et al*., 2024), which is particularly interesting given the different forms of tetraploidy. It is anticipated that more cases will be uncovered as high-quality reference genomes become available.

Key challenges of aquaculture include successful fertilization of eggs, healthy development and raising healthy fry in captivity (Teletchea, 2021). *Brd2* is involved in reproduction, embryo development, and the differentiation of the digestive tract in fish (Rhee *et al*., 1998) (DiBenedetto *et al*. 2008) (Murphy *et al*. 2017). In humans, a variant in *brd2* is strongly associated with celiac disease, an autoimmune disorder affecting the small intestine triggered by gluten consumption (Spanish Consortium on the Genetics of Coeliac Disease (CEGEC) *et al*., 2011). Given the roles of *brd2* in developmental processes across species, these parallel selected variants may contributed developmental adaptation to aquaculture conditions. We also identified parallel selection on brain-related homeolog genes, *abr*, *brd2*. We found *Abr* and *brd2* to be highly expressed in the brain confirming a neurological role in salmonid fishes. It has been reported that *abr* has an essential role in the regulation of cerebellar development, (Chuang *et al*., 1995; Mulherkar *et al*., 2014; Guo, Yang and Shi, 2020). In aquaculture environments, fish are raised under conditions of higher density than in the wild, with controlled lighting, temperature, and salinity levels. It is hypothesized that in such artificial environments, advantageous alleles in captivity may differ from beneficial alleles in nature. Disease outbreaks are a major challenge in aquaculture, regardless of species (Cain, 2022). The immune genes on homeologous regions under selection, may contribute to higher survival rates under aquaculture conditions by enhancing resistance to infectious diseases.

### Sema genes under selection in the Western Norwegian aquaculture group

Building on understanding neural-related genetic selection in aquaculture, we underscore the sema genes (Alto and Terman, 2017). Gene ontology (GO) enrichment analysis revealed that genetic variants under selection related to semaphorins were over-represented in the Western Norwegian aquaculture group (sema 4ab, sema5ba, sema6d, sema6e). Semaphonin is involved in axon guidance and synapse formation (Luo, Raible and Raper, 1993; Jongbloets and Pasterkamp, 2014). Semaphorins have also been detected under selection in previous studies of aquaculture Atlantic salmon populations (Gutierrez, Yáñez and Davidson, 2016) (Naval-Sanchez *et al*., 2020), supporting our findings. Such genes under lineage-specific selection should also contribute to the domestication of Atlantic salmon.

It is reported that structural variation showing genetic divergence between farmed and wild Atlantic salmon is linked to synaptic genes responsible for behavioral variation in Atlantic salmon (Bertolotti *et al*., 2020). Suggesting that individuals with greater stress tolerance have been selectively favored in aquaculture environments. Meanwhile, it is reported that fast-growing fish tend to be more aggressive (Nicieza and Metcalfe, 1999). So, it is also possible that the fast-growing fish have been selected, potentially leading to the inadvertent selection of genes associated with aggressive behaviors as a byproduct.

## CONCLUSION

Our study underscores the significant role of whole genome duplication (WGD) in the rapid adaptation of Atlantic salmon in aquaculture. We detected shared selective sweeps on the same variants in MHC genes in aquaculture groups across continental lineages. This finding revealed that the interplay of long-term balancing selection and recent human-induced selection shaped the evolutionary process at the MHC locus. Additionally, we observed selection on the *abr* gene pair in the homeologous region, originating from the whole genome duplication, highlighting the contribution of WGD in maintaining genomic variation and potentially reducing pleiotropy through sub-functionalization. This unique type of “parallel” selection also contributes to adapting to the intensive artificial conditions of aquaculture. Similar patterns of parallel selection in homeologous genes, though involving different genes, have also been observed in other species, such as common carp and Brassica. This is particularly interesting, considering the different polyploidization mechanisms. These cases illustrate the broader impact of WGD on domestication. Our research emphasizes the importance of maintaining genetic diversity to support sustainable aquaculture practices.

## MATERIAL & METHODS

### Studied populations

The present study was conducted using 370 individuals from two pairs of wild and aquaculture Atlantic salmon populations from two distinct phylogeographical lineages. The purpose of this design was to perform two independent comparisons between domestic strains and local wild counterpart populations to investigate genomic regions subject to putative artificial selection and their potential functions. Paired-end, short-read whole-genome sequencing data was retrieved from multiple publicly available datasets. We used n=80 aquaculture samples originally derived from 3 different source rivers in North America (NCBI Project No. PRJEB34225), n = 112 aquaculture Norwegian samples from the MOWI strain that is derived from the River Bolstad, in the Vosso drainage, River Årøy in Sogne og Fjordane and from marine captures near Osterfjord and Sotra, all in Western Norway (Glover *et al*., 2009) (NCBI Project No. PRJEB47441). As the closest available wild counterparts, we used n = 80 wild North American individuals from 8 Quebec rivers and n = 98 wild Norwegian individuals from 17 rivers in Western Norway (NCBI Project No. PRJEB38061, Bertolotti *et al*., 2020) (see **Figure 1**.).

In North America. Aquaculture samples were used from Gaspe of Quebec (Gaspe, n=16), St John River (n=53) and Penobscot River (n=11). Wild samples came from 8 rivers: Malbaie Charlevoix (n=10), Bonaventure (n=10), Petite riviere Cascapedia (n=10), Laval (n=10), De la Chaloupe (n=10), Jupiter (n=10), Du Vieux Fort (n=10) and Saint-Paul (n=10). (B) Geographic distribution of Atlantic salmon population from Western Norway. Aquaculture samples come from MOWI Norwegian breeding line (n=112). Wild counterparts were collected among 17 rivers: Vorma (Vorm, n=2), Suldalslågen (Suld, n=10), Vikedalselva i Vindafjord (Vike, n=10), Etneelva (Etne, n=2), Eidfjordvassdraget (Eidf, n=2), Oselva i Os (Osel, n=2), Loneelva i Osterøy (Lone, n=2), Flåmselva (Flam, n=10), Lærdalselva (Laer, n=2), Årøy (Aroy, n=10), Daleelva (Dale, n=2), Flekkeelva (Flek, n=2), Nausta (Naus, n=10), Jølstra (Jols, n=10), Oselvvassdraget (Osen, n=10), Ryggelva (Rygg, n=10) and Gloppenelva (Glop, n=2).

### Data preparation

Mapping, variant calling and quality control were performed independently for each of the four datasets (Aquaculture Norwegian, Wild Norwegian, Aquaculture North American, Wild North American) using the same pipeline and parameters. Illumina reads were mapped to the Atlantic salmon genome (Ssal_v3.1; GCA_905237065.2 ; Stenløkk, 2023) using SAMtools (Li *et al*., 2009). Duplicate reads were marked with gatk4-spark:4.3.0.0 MarkDuplicates. Genetic variation was identified using gatk4-spark:4.3.0.0 HaplotypeCaller with default parameters and the individual genotypes were merged with gatk4 - spark:4.3.0.0 GenotypeGVCFs. To avoid the detection of false positives, variants were filtered by the following parameters with gatk4-spark:4.3.0.0 (McKenna *et al*., 2010) : « QD < 2.0 » which filters out variants with a low quality score by depth of coverage; « QUAL < 50.0 » to sort variants with a low-quality score; « SOR > 4.0 » that removes variants where the number of reads supporting the reference allele is significantly skewed towards one strand (forward or reverse) compared to the other; « FS > 60.0 » to filter out variants whose reads supporting the reference allele versus the alternative allele are significantly different between the two strands; and finally « ReadPosRankSum < -8.0 » to remove variants with a low score for the difference between the mean position of the reference allele and the alternative allele among the reads supporting the variant (https://gatk.broadinstitute.org/hc/en-us). From this large pruning, we kept only biallelic markers (SNPs) with depth greater than 4 and less than 30, confident call rate greater than 90% (GQ) and missingness lower than 10% using VCFtools (Danecek *et al*., 2011).

### Population structure analysis

We used two clustering methods to investigate genetic structure among populations and have a broad representation of the genetic differentiation between North American and West Norwegian lineages. Further filtering removed loci deviating from Hardy–Weinberg equilibrium, loci with minor allele frequency (MAF) lower than 5% and linked SNPs (i.e. 50 SNP windows, 10 SNP sliding windows and an R^2^ threshold of 0.5) using PLINK v1.9 (Purcell *et al*., 2007). We ran ADMIXTURE (Alexander, Novembre and Lange, 2009; Alexander and Lange, 2011) with a number of ancestral populations set from 1 to 10 (K). The optimal K was selected based on the lowest cross-validation error. We then conducted a Principal Component Analysis (PCA) using PLINK v1.9 (Purcell *et al*., 2007) on the same filtered dataset. The results of both analyses (i.e eigenvectors and genetic variation from PCA; genetic components from ADMIXTURE) were visualized in R 4.2.2 using the *ggplot2* (Wickham, 2011) and *pophelper* (Francis, 2017) R packages.

### Detection of selection sweeps

For the detection of putative genomic regions under artificial selection in domestic strains, we applied a genome scan using three statistical methods based on (1) population differentiation (F_ST_) (Weir and Cockerham, 1984) ; (2) site frequency (Tajima’s D) (Tajima, 1989) and (3) cross population test of extended haplotype homozygosity (XP-EHH) (Sabeti *et al*., 2002). These tests were first carried out to compare aquaculture and wild populations from the same geographical region independently in order to detect lineage-specific selection signatures. Then, we investigated whether some genomic regions putatively under selection are shared by populations of different origins, indicative of parallel evolution. In the context of domestication, we expect two main scenarios in terms of selective sweeps. A hard selective sweep occurs when a beneficial allele arises and rapidly reaches fixation within a population. There is little genetic variation surrounding the selected allele because the advantageous variant quickly “sweeps” through the population, leaving little time for new mutations or recombination to occur. As a result, the genetic diversity around the selected site is reduced, and the favored allele is present on a single haplotype in the population. A soft selective sweep occurs when multiple beneficial alleles, either derived from different mutations or standing genetic variation, rise in frequency and are favored within a population. During a soft sweep, there can be multiple haplotypes carrying different advantageous alleles and the genetic diversity around the selected site is not as reduced as in a hard sweep, and multiple haplotypes may show evidence of positive selection.

First, F_ST_ identifies selection using differences in allele frequencies between populations, here between domesticated and wild populations from each geographic area independently. These statistics were calculated according to the pairwise Weir and Cockerham’s (1984) estimator using VCFtools (Danecek *et al*., 2011) in 5,000 bp non-overlapping sliding windows using wild individuals as the reference population. We extracted the top 1% values as outliers. As F_ST_ allows the detection of large changes in allele frequency, the test is particularly sensitive to hard sweeps, which involve strong selection. However, when using the sliding window approach, F_ST_ values are primarily influenced by the surrounding haplotype. Here the window size is typically small enough that the background haplotype is not a critical factor and indicates that F_ST_ values are unlikely to distinguish between hard and soft sweeps. The strength of selection will matter and it may be problematic to differentiate it from genetic drift following a founder bottleneck and small effective population size of captivity using this test.

Secondly, Tajima’s D (Tajima’s D) assumes that neutral genetic variation is maintained by a balance between mutation, genetic drift, and gene flow. The test is computed as the ratio between the number of segregating sites and the average pairwise nucleotide differences. Deviations from this neutral evolution can indicate putative selection or demographic events. Markers under selection can be expressed as positive and negative values, while zero values indicate a putative absence of selection. A significant positive Tajima’s D value may indicate an excess of intermediate-frequency alleles, which could be caused by balancing selection, population subdivision, or recent population expansion. A significant negative Tajima’s D value may indicate an excess of low-frequency alleles, which could be caused by positive selection, population bottlenecks, or purifying selection. In our study, Tajima’s D was conducted on the two aquaculture populations independently by non-overlapping sliding windows of 20,000 bp. As the domestication involved bottlenecks and focus on directional selection, we are more interested in negative values. The lowest values at the 1% level were taken as the significance level.

Finally, we investigated patterns of extended haplotypes between pairs of populations using the *rehh* R package (Gautier, Klassmann and Vitalis, 2017). Haplotype analyses are based on extended haplotype homozygosity (EHH) that represents the model of a selective sweep in which an adaptive de novo mutation appears on a haplotype and rapidly evolves towards fixation, thus reducing diversity around the locus (Sabeti *et al*., 2002). A hard sweep induces a signal of high haplotype homozygosity to be observed extending from an adaptive locus as strong rapid selection causes strong linkage that is not broken by recurrent mutation or recombination. In other words, under conditions of positive selection, a beneficial new allele can rapidly increase in frequency and create long stretches of homozygosity, or identical copies of the haplotype, around the selected site. This process results in longer and linked haplotypes detectable by EHH methods. In our case, we applied a cross-population (XP-EHH) test that compares the distribution of haplotype homozygosity between aquaculture and wild populations. When there is divergent selection between the two environments (wild and aquaculture), the allele selected in the focal population (farm) will be on a longer haplotype. This signal will not be present when selection is concordant between the two environments. Negative scores of XP-EHH suggest that selection occurred in the reference population, whereas positive scores show selection in the domestic population. Unlike Tajima and F_ST_, these tests cannot directly handle SNP genotypes because they require haplotypes. Added to the first filtering (see *Data preparation* section), variants with Minor Allele Frequency < 0.05 were removed. The remaining variants were phased to infer the haplotypes of each population using BEAGLE v4.1 (Browning and Browning, 2007) with a window size of 10,000 bp and 600 bp overlap. One-sided p-values were obtained as −log10(1-Φ(XP-EHH)), where Φ(XP-EHH) represents the Gaussian distribution function for the statistic. False Discovery Rate (FDR) test was performed to estimate the number of false positives and determine the significant threshold. We used −log10(p-value) = 4 (alpha = 0,0001) as a threshold to define significant values of XP-EHH. As XP-EHH searches for the extended haplotype, which is typical of a hard sweep, this method will be more sensitive in detecting hard sweeps than soft selective sweeps. In soft sweeps, the selected allele may have arisen and segregated as neutral variation in the wild, resulting in it being found in multiple haplotypes and the weaker signals extended haplotypes may be difficult to detect using XP-EHH, depending on the allele frequency at the time of domestication (Innan and Kim, 2004). Nevertheless, the combination of the three methods is particularly promising to increase the probability of detecting true directional selection signals.

Tests for lineage-specific and parallel genomic patterns arising from domestication and artificial selection were carried out using three genome-wide detection methods, based on site-frequency (Tajima’s D, Tajima, 1989), population differentiation (F_ST_, Weir and Cockerham, 1984) and cross population extended haplotype homozygosity patterns (XP-EHH, Sabeti *et al*., 2002). Selective sweeps during domestication and positive selection can be categorized into hard and soft sweeps. In a hard sweep, a new beneficial allele rapidly increases in frequency, reducing genetic variation in its vicinity and so appearing on a single haplotype; whereas a soft sweep involves multiple haplotypes that each contain the selected allele spreading through the population, as a result of recurrent mutation or because it is selected from standing genetic variation, with more genetic diversity around the selected site. Tajima’s D, F_ST_ and XP-EHH are especially sensitive to detecting hard sweeps, but the detection of soft sweeps is still possible. The analyses were carried out on 1,139,312 SNPs for (Tajima’s D) and F_ST_ and 1,053,324 SNPs for XP-EHH, depending on the filtering requirements.

### SNP annotation

Significant windows or SNPs from the three comparative analyses were intersected using BEDtools (Quinlan, 2014) to examine markers detected by multiple methods and thus increase the repeatability of the results. For each geographical region, three intersections were tested: (i) F_ST_/Tajima’s D, (ii) F_ST_/XP-EHH, (iii) Tajima’s D/XP-EHH. Candidate variants detected by two or more methods were then intersected with annotated genes in the Atlantic salmon reference genome (Ssal_v3.1; GCA_905237065.2 ; Stenløkk, 2023) using BEDtools window to identify variants under selection that are contained in the same windows as the genes or that are directly mapped to one or more genes.

The overlapping genes (Patro *et al*., 2017) were input into ShinyGO 0.77 (Ge, Jung and Yao, 2020); http://bioinformatics.sdstate.edu/go/) to extract biological functions and enrichment. All the intersects were tested independently to observe population-specific signatures and the F_ST_/XP-EHH intersect was later used to test for parallel evolution as it shared the most genomic regions detected under selection. We then tested parallel evolution using only the significant SNPs from XP-EHH, as this was the most suitable method.

### Gene expression analysis

We used available RNA sequencing data online 1) from early development stages (AQUA-FAANG project No. PRJEB51855) and 2) from tissues (NCBI Project No. SRP011583) including brain, eye, gill, gut, head kidney, heart, kidney, liver, muscle, nose, ovary, caecum, skin, spleen and testis. Data were quantified using the latest version of the Ensembl transcriptome (Ssa v.3.1). The fastq files were first filtered using fastp v0.23.2 (Chen, 2023) with default settings in order to remove potential sequencing adaptors in the reads. Then, the reads were mapped using the lightweight mapper salmon v1.5.2 (Patro *et al*., 2017) which tracks, by default, the position and orientation of all mapped fragments. This information is used in conjunction with the abundances from online inference to compute per-fragment conditional probabilities (Patro *et al*., 2017). Raw-counted data was then visualized using the package *ggplot* in R.

### Test for balancing selection on the MHC region on chromosome 14

Investigation of balancing selection was carried out using Tajima’s D (Tajima, 1989). Tajima’s D value was computed in the MHC region on chromosome 14 between 64320000 and 64340000 bp. Then, Tajima’s D was computed on 500 windows of 20kb with more than 15 SNPs, randomly selected across the genome. We used variant effect predictor (McLaren *et al*., 2016) to predict SNPs consequences on coding genes.

## ACKNOWLEDGEMENTS

The authors thank Joris Bertrand from the Laboratoire Génome et Développement des Plantes (LGDP) in Perpignan (France) for his advice on genetic structure analysis. PB was supported by EUR TULIP-GSR N°ANR-18-EUR-0019 a grant managed by the French National Research Agency (ANR) in the framework of the program “Investing for the FutureThe study was supported by The Research Council of Norway (grant nos. 325874, 275310 and 221734). We acknowledge the use of the Orion computing cluster at the Norwegian University of Life Sciences (NMBU). We thank Louise Chavarie for her comments on the geography of Canada.

## DATA AVAILABILITY

Script used for the analyses in this work are available at https://github.com/paulinebuso/ATLANTIDES_pipelines

## References

Alberto, F.J. et al. (2018) ‘Convergent genomic signatures of domestication in sheep and goats’, Nature Communications, 9(1), p. 813. Available at: 10.1038/s41467-018-03206-y.

Alexander, D.H. and Lange, K. (2011) ‘Enhancements to the ADMIXTURE algorithm for individual ancestry estimation’, BMC Bioinformatics, 12(1), p. 246. Available at: 10.1186/1471-2105-12-246.

Alexander, D.H., Novembre, J. and Lange, K. (2009) ‘Fast model-based estimation of ancestry in unrelated individuals’, Genome Research, 19(9), pp. 1655–1664. Available at: 10.1101/gr.094052.109.

Alto, L.T. and Terman, J.R. (2017) ‘Semaphorins and their Signaling Mechanisms’, in J.R. Terman (ed.) Semaphorin Signaling. New York, NY: Springer New York (Methods in Molecular Biology), pp. 1–25. Available at: 10.1007/978-1-4939-6448-2_1.

Amill, F., et al. (2024) ‘Characterization of gill bacterial microbiota in wild Arctic char ( *Salvelinus alpinus* ) across lakes, rivers, and bays in the Canadian Arctic ecosystems’, Microbiology Spectrum. Edited by K.A. Kormas, 12(3), pp. e02943–23. Available at: 10.1128/spectrum.02943-23.

Arima, K. et al. (2011) ‘Proteasome assembly defect due to a proteasome subunit beta type 8 (PSMB8) mutation causes the autoinflammatory disorder, Nakajo-Nishimura syndrome’, Proceedings of the National Academy of Sciences, 108(36), pp. 14914–14919. Available at: 10.1073/pnas.1106015108.

Baduel, P. et al. (2019) ‘Relaxed purifying selection in autopolyploids drives transposable element over-accumulation which provides variants for local adaptation’, Nature Communications, 10(1), p. 5818. Available at: 10.1038/s41467-019-13730-0.

Bailey, S.F. et al. (2017) ‘What drives parallel evolution?: How population size and mutational variation contribute to repeated evolution’, BioEssays, 39(1), p. e201600176. Available at: 10.1002/bies.201600176.

Bernatchez, L. and Wilson, C.C. (1998) ‘Comparative phylogeography of Nearctic and Palearctic fishes’, Molecular Ecology, 7(4), pp. 431–452. Available at: 10.1046/j.1365-294x.1998.00319.x.

Bertolotti, A.C. et al. (2020) ‘The structural variation landscape in 492 Atlantic salmon genomes’, Nature Communications, 11(1), p. 5176. Available at: 10.1038/s41467-020-18972-x.

Bicskei, B. et al. (2014) ‘A comparison of gene transcription profiles of domesticated and wild Atlantic salmon (Salmo salar L.) at early life stages, reared under controlled conditions’, BMC Genomics, 15(1), p. 884. Available at: 10.1186/1471-2164-15-884.

Bourret, V. et al. (2013) ‘SNP-array reveals genome-wide patterns of geographical and potential adaptive divergence across the natural range of A tlantic salmon ( *S almo salar* )’, Molecular Ecology, 22(3), pp. 532–551. Available at: 10.1111/mec.12003.

Bradbury, I.R. et al. (2022) ‘Genomic evidence of recent European introgression into North American farmed and wild Atlantic salmon’, Evolutionary Applications, 15(9), pp. 1436–1448. Available at: 10.1111/eva.13454.

Brawand, D. et al. (2014) ‘The genomic substrate for adaptive radiation in African cichlid fish’, Nature, 513(7518), pp. 375–381. Available at: 10.1038/nature13726.

Brenna-Hansen, S. et al. (2012) ‘Chromosomal differences between European and North American Atlantic salmon discovered by linkage mapping and supported by fluorescence in situ hybridization analysis’, BMC Genomics, 13(1), p. 432. Available at: 10.1186/1471-2164-13-432.

Browning, S.R. and Browning, B.L. (2007) ‘Rapid and Accurate Haplotype Phasing and Missing-Data Inference for Whole-Genome Association Studies By Use of Localized Haplotype Clustering’, The American Journal of Human Genetics, 81(5), pp. 1084–1097. Available at: 10.1086/521987.

Bull, J.K. et al. (2022) ‘Environment and genotype predict the genomic nature of domestication of salmonids as revealed by gene expression’, Proceedings of the Royal Society B: Biological Sciences, 289(1988), p. 20222124. Available at: 10.1098/rspb.2022.2124.

Cain, K. (2022) ‘The many challenges of disease management in aquaculture’, Journal of the World Aquaculture Society, 53(6), pp. 1080–1083. Available at: 10.1111/jwas.12936.

Chen, S. (2023) ‘Ultrafast one-pass FASTQ data preprocessing, quality control, and deduplication using fastp’, iMeta, 2(2), p. e107. Available at: 10.1002/imt2.107.

Chen, Z. et al. (2019) ‘De novo assembly of the goldfish ( *Carassius auratus* ) genome and the evolution of genes after whole-genome duplication’, Science Advances, 5(6), p. eaav0547. Available at: 10.1126/sciadv.aav0547.

Cheng, F. et al. (2016) ‘Genome resequencing and comparative variome analysis in a Brassica rapa and Brassica oleracea collection’, Scientific Data, 3(1), p. 160119. Available at: 10.1038/sdata.2016.119.

Chuang, T.H. et al. (1995) ‘Abr and Bcr are multifunctional regulators of the Rho GTP-binding protein family.’, Proceedings of the National Academy of Sciences, 92(22), pp. 10282–10286. Available at: 10.1073/pnas.92.22.10282.

Cross, T.F. and King, J. (1983) ‘Genetic effects of hatchery rearing in Atlantic salmon’, Aquaculture, 33(1–4), pp. 33–40. Available at: 10.1016/0044-8486(83)90384-8.

Danecek, P. et al. (2011) ‘The variant call format and VCFtools’, Bioinformatics, 27(15), pp. 2156– 2158. Available at: 10.1093/bioinformatics/btr330.

Darwin, C. (1868). The variation of animals and plants under domestication (Vol. 2). John murray.

Debes, P.V. et al. (2012) ‘Differences in transcription levels among wild, domesticated, and hybrid Atlantic salmon ( *Salmo salar* ) from two environments’, Molecular Ecology, 21(11), pp. 2574– 2587. Available at: 10.1111/j.1365-294X.2012.05567.x.

Détrée, C. and Gonçalves, A.T. (2019) ‘Transcriptome mining of apoptotic mechanisms in response to density and functional diets in Oncorhynchus mykiss and role in homeostatic regulation’, Comparative Biochemistry and Physiology Part D: Genomics and Proteomics, 31, p. 100595. Available at: 10.1016/j.cbd.2019.100595.

Diamond, J. (2002) ‘Evolution, consequences and future of plant and animal domestication’, Nature, 418(6898), pp. 700–707. Available at: 10.1038/nature01019.

DiBenedetto, A.J. et al. (2008) ‘Zebrafish brd2a and brd2bare paralogous members of the bromodomain-ET (BET) family of transcriptional coregulators that show structural and expression divergence’, BMC Developmental Biology, 8(1), p. 39. Available at: 10.1186/1471-213X-8-39.

Du, K. et al. (2020) ‘The sterlet sturgeon genome sequence and the mechanisms of segmental rediploidization’, Nature Ecology & Evolution, 4(6), pp. 841–852. Available at: 10.1038/s41559-020-1166-x.

Ebadi, M. et al. (2023) ‘The duplication of genomes and genetic networks and its potential for evolutionary adaptation and survival during environmental turmoil’, Proceedings of the National Academy of Sciences, 120(41), p. e2307289120. Available at: 10.1073/pnas.2307289120.

Eltaher, S. et al. (2021) ‘GWAS revealed effect of genotype × environment interactions for grain yield of Nebraska winter wheat’, BMC Genomics, 22(1), p. 2. Available at: 10.1186/s12864-020-07308-0.

Fernandez, A.R. et al. (2021) ‘Intentional and unintentional selection during plant domestication: herbivore damage, plant defensive traits and nutritional quality of fruit and seed crops’, New Phytologist, 231(4), pp. 1586–1598. Available at: 10.1111/nph.17452.

Ford, J.S. and Myers, R.A. (2008) ‘A Global Assessment of Salmon Aquaculture Impacts on Wild Salmonids’, PLoS Biology. Edited by C. Roberts, 6(2), p. e33. Available at: 10.1371/journal.pbio.0060033.

Francis, R.M. (2017) ‘POPHELPER : an R package and web app to analyse and visualize population structure’, Molecular Ecology Resources, 17(1), pp. 27–32. Available at: 10.1111/1755-0998.12509.

Frantz, L.A.F. et al. (2020) ‘Animal domestication in the era of ancient genomics’, Nature Reviews Genetics, 21(8), pp. 449–460. Available at: 10.1038/s41576-020-0225-0.

Freebern, E. et al. (2020) ‘GWAS and fine-mapping of livability and six disease traits in Holstein cattle’, BMC Genomics, 21(1), p. 41. Available at: 10.1186/s12864-020-6461-z.

Garbi, N. et al. (2003) ‘A major role for tapasin as a stabilizer of the TAP peptide transporter and consequences for MHC class I expression’, European Journal of Immunology, 33(1), pp. 264–273. Available at: 10.1002/immu.200390029.

Gautier, M., Klassmann, A. and Vitalis, R. (2017) ‘REHH 2.0: a reimplementation of the R package REHH to detect positive selection from haplotype structure’, Molecular Ecology Resources, 17(1), pp. 78–90. Available at: 10.1111/1755-0998.12634.

Ge, S.X., Jung, D. and Yao, R. (2020) ‘ShinyGO: a graphical gene-set enrichment tool for animals and plants’, Bioinformatics. Edited by A. Valencia, 36(8), pp. 2628–2629. Available at: 10.1093/bioinformatics/btz931.

Gillard, G.B. et al. (2021) ‘Comparative regulomics supports pervasive selection on gene dosage following whole genome duplication’, Genome Biology, 22(1), p. 103. Available at: 10.1186/s13059-021-02323-0.

Gjedrem, T., Gjøen, H.M. and Gjerde, B. (1991) ‘Genetic origin of Norwegian farmed Atlantic salmon’, Aquaculture, 98(1–3), pp. 41–50. Available at: 10.1016/0044-8486(91)90369-I.

Glover, K.A. et al. (2009) ‘A comparison of farmed, wild and hybrid Atlantic salmon (Salmo salar L.) reared under farming conditions’, Aquaculture, 286(3–4), pp. 203–210. Available at: 10.1016/j.aquaculture.2008.09.023.

Glover, K.A. et al. (2017a) ‘Half a century of genetic interaction between farmed and wild Atlantic salmon: Status of knowledge and unanswered questions’, Fish and Fisheries, 18(5), pp. 890–927. Available at: 10.1111/faf.12214.

Glover, K.A. et al. (2017b) ‘Half a century of genetic interaction between farmed and wild Atlantic salmon: Status of knowledge and unanswered questions’, Fish and Fisheries, 18(5), pp. 890–927. Available at: 10.1111/faf.12214.

Grimholt, U. (2018) ‘Whole genome duplications have provided teleosts with many roads to peptide loaded MHC class I molecules’, BMC Evolutionary Biology, 18(1), p. 25. Available at: 10.1186/s12862-018-1138-9.

Gundappa, M.K., et al. (2022) ‘Genome-Wide Reconstruction of Rediploidization Following Autopolyploidization across One Hundred Million Years of Salmonid Evolution’, Molecular Biology and Evolution. Edited by M. O’Connell, 39(1), p. msab310. Available at: 10.1093/molbev/msab310.

Guo, D., Yang, X. and Shi, L. (2020) ‘Rho GTPase Regulators and Effectors in Autism Spectrum Disorders: Animal Models and Insights for Therapeutics’, Cells, 9(4), p. 835. Available at: 10.3390/cells9040835.

Gutierrez, A.P., Yáñez, J.M. and Davidson, W.S. (2016) ‘Evidence of recent signatures of selection during domestication in an Atlantic salmon population’, Marine Genomics, 26, pp. 41–50. Available at: 10.1016/j.margen.2015.12.007.

Han, K. et al. (2018) ‘QTL mapping and GWAS reveal candidate genes controlling capsaicinoid content in *Capsicum*’, Plant Biotechnology Journal, 16(9), pp. 1546–1558. Available at: 10.1111/pbi.12894.

Harvey, A.C. et al. (2018) ‘Implications for introgression: has selection for fast growth altered the size threshold for precocious male maturation in domesticated Atlantic salmon?’, BMC Evolutionary Biology, 18(1), p. 188. Available at: 10.1186/s12862-018-1294-y.

Hayes, B.J., Lewin, H.A. and Goddard, M.E. (2013) ‘The future of livestock breeding: genomic selection for efficiency, reduced emissions intensity, and adaptation’, Trends in Genetics, 29(4), pp. 206–214. Available at: 10.1016/j.tig.2012.11.009.

Hewitt, G. (2000) ‘The genetic legacy of the Quaternary ice ages’, Nature, 405(6789), pp. 907–913. Available at: 10.1038/35016000.

Horn, S.S. et al. (2020) ‘GWAS identifies genetic variants associated with omega-3 fatty acid composition of Atlantic salmon fillets’, Aquaculture, 514, p. 734494. Available at: 10.1016/j.aquaculture.2019.734494.

Houston, R.D. and Macqueen, D.J. (2019) ‘Atlantic salmon ( *Salmo salar* L.) genetics in the 21st century: taking leaps forward in aquaculture and biological understanding’, Animal Genetics, 50(1), pp. 3–14. Available at: 10.1111/age.12748.

Huang, X. et al. (2022) ‘The integrated genomics of crop domestication and breeding’, Cell, 185(15), pp. 2828–2839. Available at: 10.1016/j.cell.2022.04.036.

Innan, H. and Kim, Y. (2004) ‘Pattern of polymorphism after strong artificial selection in a domestication event’, Proceedings of the National Academy of Sciences, 101(29), pp. 10667– 10672. Available at: 10.1073/pnas.0401720101.

Izawa, T. (2022) ‘Reloading DNA History in Rice Domestication’, Plant and Cell Physiology, 63(11), pp. 1529–1539. Available at: 10.1093/pcp/pcac073.

Jin, Y. et al. (2020) ‘Comparative transcriptomics reveals domestication-associated features of Atlantic salmon lipid metabolism’, Molecular Ecology, 29(10), pp. 1860–1872. Available at: 10.1111/mec.15446.

Jongbloets, B.C. and Pasterkamp, R.J. (2014) ‘Semaphorin signalling during development’, Development, 141(17), pp. 3292–3297. Available at: 10.1242/dev.105544.

Khelifi, A.F., D’Alcontres, M.S. and Salomoni, P. (2005) ‘Daxx is required for stress-induced cell death and JNK activation’, Cell Death & Differentiation, 12(7), pp. 724–733. Available at: 10.1038/sj.cdd.4401559.

Klemetsen, A. et al. (2003) ‘Atlantic salmon *Salmo salar* L., brown trout *Salmo trutta* L. and Arctic charr *Salvelinus alpinus* (L.): a review of aspects of their life histories’, Ecology of Freshwater Fish, 12(1), pp. 1–59. Available at: 10.1034/j.1600-0633.2003.00010.x.

Krkosek, M. (2010) ‘Host density thresholds and disease control for fisheries and aquaculture’, Aquaculture Environment Interactions, 1(1), pp. 21–32. Available at: 10.3354/aei0004.

Larson, G. et al. (2014) ‘Current perspectives and the future of domestication studies’, Proceedings of the National Academy of Sciences, 111(17), pp. 6139–6146. Available at: 10.1073/pnas.1323964111.

Lehner, P.J., Surman, M.J. and Cresswell, P. (1998) ‘Soluble Tapasin Restores MHC Class I Expression and Function in the Tapasin-Negative Cell Line .220’, Immunity, 8(2), pp. 221–231. Available at: 10.1016/S1074-7613(00)80474-4.

Lehnert, S.J. et al. (2018) Chromosome polymorphisms track trans-Atlantic divergence, admixture and adaptive evolution in salmon. preprint. Genomics. Available at: 10.1101/351338.

Li, H. et al. (2009) ‘The Sequence Alignment/Map format and SAMtools’, Bioinformatics, 25(16), pp. 2078–2079. Available at: 10.1093/bioinformatics/btp352.

Li, N. et al. (2018) ‘Identification of novel genes significantly affecting growth in catfish through GWAS analysis’, Molecular Genetics and Genomics, 293(3), pp. 587–599. Available at: 10.1007/s00438-017-1406-1.

Lien, S. et al. (2016) ‘The Atlantic salmon genome provides insights into rediploidization’, Nature, 533(7602), pp. 200–205. Available at: 10.1038/nature17164.

Liu, Z. et al. (2019) ‘Genome-wide association analysis of egg production performance in chickens across the whole laying period’, BMC Genetics, 20(1), p. 67. Available at: 10.1186/s12863-019-0771-7.

López, M.E. et al. (2019) ‘Comparing genomic signatures of domestication in two Atlantic salmon (*Salmo salar* L.) populations with different geographical origins’, Evolutionary Applications, 12(1), pp. 137–156. Available at: 10.1111/eva.12689.

Lukacs, M.F. et al. (2007) ‘Genomic organization of duplicated major histocompatibility complex class I regions in Atlantic salmon (Salmo salar)’, BMC Genomics, 8(1), p. 251. Available at: 10.1186/1471-2164-8-251.

Lukacs, M.F. et al. (2010) ‘Comprehensive analysis of MHC class I genes from the U-, S-, and Z-lineages in Atlantic salmon’, BMC Genomics, 11(1), p. 154. Available at: 10.1186/1471-2164-11-154.

Luo, Y., Raible, D. and Raper, J.A. (1993) ‘Collapsin: A protein in brain that induces the collapse and paralysis of neuronal growth cones’, Cell, 75(2), pp. 217–227. Available at: 10.1016/0092-8674(93)80064-L.

Marchant, A. et al. (2019) ‘The role of structural pleiotropy and regulatory evolution in the retention of heteromers of paralogs’, eLife, 8, p. e46754. Available at: 10.7554/eLife.46754.

McConnell, S.C. et al. (2016) ‘Alternative haplotypes of antigen processing genes in zebrafish diverged early in vertebrate evolution’, Proceedings of the National Academy of Sciences, 113(34). Available at: 10.1073/pnas.1607602113.

McKenna, A. et al. (2010) ‘The Genome Analysis Toolkit: A MapReduce framework for analyzing next-generation DNA sequencing data’, Genome Research, 20(9), pp. 1297–1303. Available at: 10.1101/gr.107524.110.

McLaren, W. et al. (2016) ‘The Ensembl Variant Effect Predictor’, Genome Biology, 17(1), p. 122. Available at: 10.1186/s13059-016-0974-4.

Michaelson, J.S. et al. (1999) ‘Loss of Daxx, a promiscuously interacting protein, results in extensive apoptosis in early mouse development’, Genes & Development, 13(15), pp. 1918–1923. Available at: 10.1101/gad.13.15.1918.

Mignon-Grasteau, S. et al. (2005) ‘Genetics of adaptation and domestication in livestock’, Livestock Production Science, 93(1), pp. 3–14. Available at: 10.1016/j.livprodsci.2004.11.001.

Mulherkar, S. et al. (2014) ‘The small GTPases RhoA and Rac1 regulate cerebellar development by controlling cell morphogenesis, migration and foliation’, Developmental Biology, 394(1), pp. 39–53. Available at: 10.1016/j.ydbio.2014.08.004.

Murphy, T. et al. (2017) ‘Knockdown of epigenetic transcriptional co-regulator Brd2a disrupts apoptosis and proper formation of hindbrain and midbrain-hindbrain boundary (MHB) region in zebrafish’, Mechanisms of Development, 146, pp. 10–30. Available at: 10.1016/j.mod.2017.05.003.

Nakajima, T. et al. (2019) ‘Common carp aquaculture in Neolithic China dates back 8,000 years’, Nature Ecology & Evolution, 3(10), pp. 1415–1418. Available at: 10.1038/s41559-019-0974-3.

Naval-Sanchez, M. et al. (2020) ‘Changed Patterns of Genomic Variation Following Recent Domestication: Selection Sweeps in Farmed Atlantic Salmon’, Frontiers in Genetics, 11, p. 264. Available at: 10.3389/fgene.2020.00264.

Nicieza, A.G. and Metcalfe, N.B. (1999) ‘Costs of rapid growth: the risk of aggression is higher for fast-growing salmon’, Functional Ecology, 13(6), pp. 793–800. Available at: 10.1046/j.1365-2435.1999.00371.x.

Nugent, C.M. et al. (2024) ‘Post-glacial recolonization and multiple scales of secondary contact contribute to contemporary Atlantic salmon ( *Salmo salar* ) genomic variation in North America’, Journal of Biogeography, p. jbi.14852. Available at: 10.1111/jbi.14852.

Oleksyk, T.K., Smith, M.W. and O’Brien, S.J. (2010) ‘Genome-wide scans for footprints of natural selection’, Philosophical Transactions of the Royal Society B: Biological Sciences, 365(1537), pp. 185–205. Available at: 10.1098/rstb.2009.0219.

Ortmann, B. et al. (1997) ‘A Critical Role for Tapasin in the Assembly and Function of Multimeric MHC Class I-TAP Complexes’, Science, 277(5330), pp. 1306–1309. Available at: 10.1126/science.277.5330.1306.

Pajic, P. et al. (2019) ‘Independent amylase gene copy number bursts correlate with dietary preferences in mammals’, eLife, 8, p. e44628. Available at: 10.7554/eLife.44628.

Patro, R. et al. (2017) ‘Salmon provides fast and bias-aware quantification of transcript expression’, Nature Methods, 14(4), pp. 417–419. Available at: 10.1038/nmeth.4197.

Purcell, S. et al. (2007) ‘PLINK: A Tool Set for Whole-Genome Association and Population-Based Linkage Analyses’, The American Journal of Human Genetics, 81(3), pp. 559–575. Available at: 10.1086/519795.

Quinlan, A.R. (2014) ‘BEDTools: The Swiss-Army Tool for Genome Feature Analysis’, Current Protocols in Bioinformatics, 47(1). Available at: 10.1002/0471250953.bi1112s47.

Redmond, A.K. et al. (2023) ‘Independent rediploidization masks shared whole genome duplication in the sturgeon-paddlefish ancestor’, Nature Communications, 14(1), p. 2879. Available at: 10.1038/s41467-023-38714-z.

Rhee, K. et al. (1998) ‘Expression and potential role of *Fsrg1*, a murine bromodomain-containing homologue of the *Drosophila* gene *female sterile homeotic*’, Journal of Cell Science, 111(23), pp. 3541–3550. Available at: 10.1242/jcs.111.23.3541.

Robertson, F.M. et al. (2017) ‘Lineage-specific rediploidization is a mechanism to explain time-lags between genome duplication and evolutionary diversification’, Genome Biology, 18(1), p. 111. Available at: 10.1186/s13059-017-1241-z.

Robinson, J.T. et al. (2011) ‘Integrative genomics viewer’, Nature Biotechnology, 29(1), pp. 24–26. Available at: 10.1038/nbt.1754.

Rondeau, E.B., et al. (2023) ‘Insights from a chum salmon ( *Oncorhynchus keta* ) genome assembly regarding whole-genome duplication and nucleotide variation influencing gene function’, G3: Genes, Genomes, Genetics. Edited by D.-J. De Koning, 13(8), p. jkad127. Available at: 10.1093/g3journal/jkad127.

Rougemont, Q. and Bernatchez, L. (2018) ‘The demographic history of Atlantic salmon ( *Salmo salar* ) across its distribution range reconstructed from approximate Bayesian computations*: ATLANTIC SALMON HISTORY AND LINKED SELECTION’, Evolution, 72(6), pp. 1261–1277. Available at: 10.1111/evo.13486.

Sabeti, P.C. et al. (2002) ‘Detecting recent positive selection in the human genome from haplotype structure’, Nature, 419(6909), pp. 832–837. Available at: 10.1038/nature01140.

Salman-Minkov, A., Sabath, N. and Mayrose, I. (2016) ‘Whole-genome duplication as a key factor in crop domestication’, Nature Plants, 2(8), p. 16115. Available at: 10.1038/nplants.2016.115.

Sever, L. et al. (2014) ‘Expression of tapasin in rainbow trout tissues and cell lines and up regulation in a monocyte/macrophage cell line (RTS11) by a viral mimic and viral infection’, Developmental & Comparative Immunology, 44(1), pp. 86–93. Available at: 10.1016/j.dci.2013.11.019.

Sokolkova, A. et al. (2020) ‘Genome-wide association study in accessions of the mini-core collection of mungbean (Vigna radiata) from the World Vegetable Gene Bank (Taiwan)’, BMC Plant Biology, 20(S1), p. 363. Available at: 10.1186/s12870-020-02579-x.

Solberg, M.F. et al. (2020) ‘Domestication leads to increased predation susceptibility’, Scientific Reports, 10(1), p. 1929. Available at: 10.1038/s41598-020-58661-9.

Spanish Consortium on the Genetics of Coeliac Disease (CEGEC) et al. (2011) ‘Dense genotyping identifies and localizes multiple common and rare variant association signals in celiac disease’, Nature Genetics, 43(12), pp. 1193–1201. Available at: 10.1038/ng.998.

Spurgin, L.G. and Richardson, D.S. (2010) ‘How pathogens drive genetic diversity: MHC, mechanisms and misunderstandings’, Proceedings of the Royal Society B: Biological Sciences, 277(1684), pp. 979–988. Available at: 10.1098/rspb.2009.2084.

Stenløkk, K. S. R. (2023). Genomic structural variations as drivers of adaptation in salmonid fishes.

Sundaram, A.Y.M. et al. (2020) ‘An Illumina approach to MHC typing of Atlantic salmon’, Immunogenetics, 72(1–2), pp. 89–100. Available at: 10.1007/s00251-019-01143-8.

Tajima, F. (1989) ‘Statistical method for testing the neutral mutation hypothesis by DNA polymorphism.’, Genetics, 123(3), pp. 585–595. Available at: 10.1093/genetics/123.3.585.

Teixeira, J.C. et al. (2015) ‘Long-Term Balancing Selection in LAD1 Maintains a Missense Trans-Species Polymorphism in Humans, Chimpanzees, and Bonobos’, Molecular Biology and Evolution, 32(5), pp. 1186–1196. Available at: 10.1093/molbev/msv007.

Teletchea, F. (2021) ‘Fish domestication in aquaculture: 10 unanswered questions’, Animal Frontiers, 11(3), pp. 87–91. Available at: 10.1093/af/vfab012.

The Brassica rapa Genome Sequencing Project Consortium et al. (2011) ‘The genome of the mesopolyploid crop species Brassica rapa’, Nature Genetics, 43(10), pp. 1035–1039. Available at: 10.1038/ng.919.

Tsukamoto, K. et al. (2012) ‘Long-Lived Dichotomous Lineages of the Proteasome Subunit Beta Type 8 (PSMB8) Gene Surviving More than 500 Million Years as Alleles or Paralogs’, Molecular Biology and Evolution, 29(10), pp. 3071–3079. Available at: 10.1093/molbev/mss113.

Veríssimo, A., et al. (2023) ‘An Ancestral Major Histocompatibility Complex Organization in Cartilaginous Fish: Reconstructing MHC Origin and Evolution’, Molecular Biology and Evolution. Edited by M. Yeager, 40(12), p. msad262. Available at: 10.1093/molbev/msad262.

Vigne, J.-D. (2011) ‘The origins of animal domestication and husbandry: A major change in the history of humanity and the biosphere’, Comptes Rendus Biologies, 334(3), pp. 171–181. Available at: 10.1016/j.crvi.2010.12.009.

Wang, B. et al. (2020) ‘Genome-wide selection and genetic improvement during modern maize breeding’, Nature Genetics, 52(6), pp. 565–571. Available at: 10.1038/s41588-020-0616-3.

Wang, G.-D. et al. (2014) ‘Domestication Genomics: Evidence from Animals’, Annual Review of Animal Biosciences, 2(1), pp. 65–84. Available at: 10.1146/annurev-animal-022513-114129.

Wang, M. et al. (2024) ‘Asymmetric and parallel subgenome selection co-shape common carp domestication’, BMC Biology, 22(1), p. 4. Available at: 10.1186/s12915-023-01806-9.

Weir, B.S. and Cockerham, C.C. (1984) ‘Estimating F-Statistics for the Analysis of Population Structure’, Evolution, 38(6), p. 1358. Available at: 10.2307/2408641.

Weir, B.S. and Cockerham, C.C. (2023) ‘Estimating F-Statistics for the Analysis of Population Structure’.

Wickham, H. (2011) ‘ggplot2: ggplot2’, Wiley Interdisciplinary Reviews: Computational Statistics, 3(2), pp. 180–185. Available at: 10.1002/wics.147.

Wiener, P. and Wilkinson, S. (2011) ‘Deciphering the genetic basis of animal domestication’, Proceedings of the Royal Society B: Biological Sciences, 278(1722), pp. 3161–3170. Available at: 10.1098/rspb.2011.1376.

Witt, K.E. and Huerta-Sánchez, E. (2019) ‘Convergent evolution in human and domesticate adaptation to high-altitude environments’, Philosophical Transactions of the Royal Society B: Biological Sciences, 374(1777), p. 20180235. Available at: 10.1098/rstb.2018.0235.

Wood, T.E., Burke, J.M. and Rieseberg, L.H. (2008) ‘Parallel genotypic adaptation: when evolution repeats itself’.

Xu, P. et al. (2019) ‘The allotetraploid origin and asymmetrical genome evolution of the common carp Cyprinus carpio’, Nature Communications, 10(1), p. 4625. Available at: 10.1038/s41467-019-12644-1.

Yáñez, J.M. et al. (2023) ‘Genome-wide association and genomic selection in aquaculture’, Reviews in Aquaculture, 15(2), pp. 645–675. Available at: 10.1111/raq.12750.

Yao, Z. et al. (2014) ‘Death Domain-associated Protein 6 (Daxx) Selectively Represses IL-6 Transcription through Histone Deacetylase 1 (HDAC1)-mediated Histone Deacetylation in Macrophages’, Journal of Biological Chemistry, 289(13), pp. 9372–9379. Available at: 10.1074/jbc.M113.533992.

Zhang, K., Wang, X. and Cheng, F. (2019) ‘Plant Polyploidy: Origin, Evolution, and Its Influence on Crop Domestication’, Horticultural Plant Journal, 5(6), pp. 231–239. Available at: 10.1016/j.hpj.2019.11.003.

Zhang, Y. et al. (2019) ‘Genetic correlation of fatty acid composition with growth, carcass, fat deposition and meat quality traits based on GWAS data in six pig populations’, Meat Science, 150, pp. 47–55. Available at: 10.1016/j.meatsci.2018.12.008.

